# Predicting Epigenomic Functions of Genetic Variants in the Context of Neurodevelopment via Deep Transfer Learning

**DOI:** 10.1101/2021.02.02.429064

**Authors:** Boqiao Lai, Sheng Qian, Hanwen Zhang, Siwei Zhang, Alena Kozlova, Jubao Duan, Xin He, Jinbo Xu

## Abstract

Decoding the regulatory effects of non-coding variants is a key challenge in understanding the mechanisms of gene regulation as well as the genetics of common diseases. Recently, deep learning models have been introduced to predict genome-wide epigenomic profiles and effects of DNA variants, in various cellular contexts, but they were often trained in cell lines or bulk tissues that may not be related to phenotypes of interest. This is particularly a challenge for neuropsychiatric disorders, since the most relevant cell and tissue types are often missing in the training data of such models.

To address this issue, we introduce a deep transfer learning framework termed MetaChrom that takes advantage of both a reference dataset - an extensive compendium of publicly available epigenomic data, and epigenomic profiles of cell types related to specific phenotypes of interest. We trained and evaluated our model on a comprehensive set of epigenomic profiles from fetal and adult brain, and cellular models representing early neurodevelopment. MetaChrom predicts these epigenomic features with much higher accuracy than previous methods, and than models without the use of reference epigenomic data for transfer learning. Using experimentally determined regulatory variants from iPS cell-derived neurons, we show that MetaChrom predicts functional variants more accurately than existing non-coding variant scoring tools. By combining genome-wide association study (GWAS) data with MetaChrom predictions, we prioritized 31 SNPs for Schizophrenia (SCZ). These candidate SNPs suggest potential risk genes of SCZ and the biological contexts where they act.

In summary, MetaChrom is a general transfer learning framework that can be applied to the study of regulatory functions of DNA sequences and variants in any disease-related cell or tissue types. The software tool is available at https://github.com/bl-2633/MetaChrom and a prediction web server is accessible at https://metachrom.ttic.edu/.

## 1 Introduction

Gene expression is controlled by the epigenome, which represents the regulatory activities of noncoding genome. Various sequencing technologies such as ATAC-seq[7], DNase-seq[70] and ChIP-seq[56] have been developed to measure the epigenomic landscapes across many cellular contexts, including histone marks, transcription factor (TF) binding and chromatin accessibility. These epigenomic annotations facilitate the study of regulatory functions of non-coding genomic regions [41, 77]. This is particularly important in the context of human disease studies, as the vast majority of disease-associated genetic variants are located in noncoding regions[12, 47]. Leveraging these resources, researchers have developed machine learning models to learn features of DNA sequences that may predict epigenomic profiles such as protein binding sites, chromatin accessibility, histone marks and methylation of DNA sequences[2, 39, 4, 58, 14, 65, 87]. Many recent methods use deep neural networks, a powerful paradigm for classification and prediction, [36, 69, 10]. In particular, Convolutional neural networks (CNNs) and related models outperform traditional methods in sequence-based prediction of protein-DNA/RNA interaction and chromatin accessibility [91, 1, 32, 38, 18, 27]. Once trained, these models can then be used to predict the likely regulatory effect of a DNA variant[91, 32] by comparing predicted epigenomic properties of different alleles. Notably, models have been trained on large publicly available epigenomic datasets such as ENCODE and Roadmap Epigenomics Consortium[12, 37], providing rich annotations of regulatory functions of variants.

Regulatory functions of DNA sequences and variants, however, are often cell type specific. Pre-trained models may not include cell types of interest and thus not able to provide the correct variant annotations in those cell types. It is thus desirable for a researcher to train specific models for the epigenomic data in the cell types of his/her interest. However, training deep learning models, from the scratch, with small training sets is challenging. While it is possible to train a single model combining all available datasets, including large public consortium data, the process of data collection, harmonization and training involves considerable efforts.

We proposed to address this challenge with a deep transfer learning framework. Transfer learning is a general machine learning approach that leverages knowledge and models gained from one domain to a related domain [74, 43]. In our case, we leverage models learned from external datasets to build a better deep neuron network for a particular dataset of interest. Specifically, our moethod is built upon Convolutional Residual Networks (ResNet)[30, 50]. ResNet is a technique that may train very deep CNNs to enhance the predictive power, and has been proven effective in computational biology problems such as RNA binding motif discovery[34] and protein folding[84]. To train the ResNet model for an epigenomic dataset from cell types of interest, our method, named MetaChrom, uses transfer learning[74, 43]. It uses a CNN-based meta-feature extractor to learn rich sequence features from the 919 external epigenomic profiles of diverse cell and tissue types from the ENCODE Project and Roadmap Epigenomics Consortium[12, 37]. Then it combines this meta-feature extractor with ResNet to learn a sequence model for the epigenomic dataset of interest. This strategy thus has the advantage of rich representation of ResNet, while avoids overfitting by using sequence features learned from external datasets.

We demonstrate the advantage of MetaChrom in the context of studying neuropsychiatric disorders. Because of the difficulty of measuring epigenomes in developing human brain, previous studies often focused on a limited set of postmortem brain samples (usually adult) [16, 21, 6]. However, adult brain samples may not capture the early stages of neurodevelopment, which are important for many neuropsychiatric disorders[16, 80, 90]. As a result, relevant cell and tissue types are under-represented in the training set of current variant analysis tools, making it difficult to annotate functions of variants in cell types related to neurodevelopment. We bridge this gap by collecting a large collection of epigenomic data in both fetal and postmortem brains, and from cellular models of early neurodevelopment, including brain organoid and induced Pluripotent Stem Cell (iPSC) derived neuronal cells[78]. Regulatory sequences in these cellular models, as our recent work demonstrated, differ substantially from those in adult brains, and are highly enriched with risk variants of neuropsychiatric traits[90, 19]. We applied MetaChrom to these datasets, while taking advantage of a large reference epigenomic dataset, to learn important DNA sequence features and predict functional effects of noncoding variants in the context of neurodevelopment and neuropsychiatric disorders. MetaChrom outperforms previous deep learning methods[91, 59] and models without transfer learning, in predicting epigenomic profiles. These higher predictive accuracy translates to better prediction of functional effects from single nucleotide variants.

We demonstrate the utility of our method in studying genetics of schizophrenia (SCZ), a complex mental disorder. The risk of SCZ has been associated with more than 100 genetic loci via genomewide-association study (GWAS), but in most loci, the causal variants remain unknown[52]. One strategy to resolve this challenge is to focus on variants with functional effects in disease relevant tissues and cell types. Such variants are more likely to be causal variants *a priori*, because functional variants are generally sparse in the human genome[28], and particularly so in a given disease context. Combining neurodevelopment-specific predictions of MetaChrom with GWAS results, we highlight 31 likely functional SNPs in 30 SCZ-associated loci. Studying these variants points to putative causal genes in these loci and the cell types and developmental stages where these variants act.

## 2 Results

### 2.1 MetaChrom: sequence-based prediction of epigenomic profiles and variant effects using transfer learning

Our goal is to train a computational model that predicts epigenomic profiles of any given DNA sequences. Our training set consists of a large number of DNA sequences, 1000 bps in length, and their functional labels, e.g. whether a sequence is in open chromatin region or not, in a given cell type. Additionally we have access to a large compendium of publicly available epigenomic profiles - the reference epigenomic data, which will be used to extract sequence features to improve model learning capability.

The modular framework we have built, MetaChrom has two major components (Fig.**??** A): (1) a meta-feature extractor (MetaFeat) pre-trained on the reference epigenomic data; (2) a ResNet based sequence encoder. The meta-feature and the encoded sequence are then combined to predict the epigenomic profiles of the sequence of interest. The meta-feature, i.e. transfer learning, component learns important sequence features from the reference set. While precise interpretation of features in deep neural networks is generally difficult, conceptually these sequence features can be viewed as certain “regulatory code”, e.g. synergistic interaction between a pair of motifs. As shown later, learning such high-level features would improve the model performance.Once a sequence-to-function model is trained, MetaChrom will be able to predict regulatory effects of genomic variants (Fig.**??** B). By comparing the predicted epigenomic profiles of two sequences differing in a single nucleotide, MetaChrom is able to assess the likely function of individual variants.

### 2.2 MetaChrom accurately predicts epigenomic profiles across neurodevelopment-related cell types

We evaluated MetaChrom in predicting epigenomic features on a collection of 31 datasets, including chromatin accessibility and histone marks of enhancers, derived from both fetal and adult brain tissues or neuronal cells (Methods, Table S1). The test sequences were obtained from chromosome 7 and 8, which were not used in the training process. For comparison, we implemented a baseline CNN model with 3 convolutional layers Such a shallow CNN model has been commonly used for predicting epigenomic profiles from DNA sequences [32, 91]. We also included DanQ in comparison, a deep hybrid recurrent neural network model developed for epigenomic prediction. Finally, we assess how well a method performs in our datasets, if it was trained on other cell types. In particular, we are interested in the question of whether average epigenomic activities of a sequence across a large collection of cell types would be a good predictor of its activity in a new cell type. Such possibility has been raised in several recent papers [63, 49]. For this analysis, we use average epigenomic profiles, from DeepSEA, across a broad range of 919 cell and tissue types/conditions.

We evaluated the model performance in predicting sequence labels in the testing data using AUPRC and AUROC, standing for Area under Precision-Recall curve (PRC) and Area under Receiver Operating Characteristic (ROC) curve, respectively. Our ResNet based transfer learning model MetaChrom achieved average AUROC = 0.89 and AUPRC = 0.51 across 31 cell-types compare to the Baseline CNN model with AUROC = 0.84 and AUPRC = 0.42 and the DanQ model with AUROC = 0.86 and AUPRC = 0.44 while the average DeepSEA baseline achieved average AUROC = 0.64 and AUPRC = 0.20 (Fig. 2). To give a detailed picture of how the models perform, we showed the Response-Operating Curves and Precision-Recall curves of the four methods in one cell type, iPS cell derived Glutamernergic neurons in Figure S1 and the complete results in Figures. S2–3. In all cell types, our model outperforms other methods.

To better understand the importance of transfer learning and the contribution of model architecture (ResNet vs. CNN) to the performance, we performed additional comparison of MetaChrom against two variants of MetaChrom: one without transfer learning and one where CNN instead of ResNet is used. When transferred knowledge is not used, our ResNet has average AUPRC=0.28 and average AUROC=0.80 across 31 cell types, compare to the baseline CNN model with AUPRC=0.39 and AUROC=0.80. The somewhat lower performance of ResNet is not surprising since ResNet is a considerably deeper network than CNN, and may suffer from overfitting and other common disadvantage of such deeper networks. When transfer leaning is used, both CNN and ResNet benefit, as shown in Fig. S4–S5, The improvement of CNN is relatively modest: with AUPRC increasing from 0.39 to 0.44 and AUROC from 0.80 to 0.835. In contrast, ResNet with the meta-feature extractor dramatically improves the performance: increasing its AUPRC from 0.28 to 0.50 and AUROC from 0.80 to 0.86.

In summary, we demonstrate that MetaChrom is a powerful framework of predicting epigenomic profiles from DNA sequences, outperforming existing methods. Its power lies in both its ResNet architecture and its ability of transfer learning from external datasets. MetaChrom offers an even larger advantage over current methods when the training data is small.

### 2.3 MetaChrom predicted functional variants are supported by evolutionary constraint and allele-specific chromatin variants

We next evaluated the accuracy of predicting functional effects of DNA variants by MetaChrom. Functionally important positions in the genome are often indicated by evolutionary constraint [26]. We thus evaluated the constraint on MetaChrom predicted variants. For all SNPs within peak regions of epigenomic data of each cell type, we computed MetaChrom scores, defined as the absolute value of the difference of MetaChrom predictions between reference and alternative alleles (Figure 1B). From these SNPs, we chose top 10,000 as predicted functional variants, and randomly sampled 100,000 variants from peak regions in the same cell type as control. We compared GERP scores, a commonly used measure of inter-species conservation [15], between functional and control SNPs. In most cell types, MetaChrom top variants have significantly higher GERP scores than random ones (Figure 3A for a subset of cell types, the rest in Figure S6), suggesting stronger evolutionary constraint. We next evaluated intra-species constraint of MetaChrom variants. Because of purifying selection, functionally deleterious variants often occur at low frequencies in the population [46, 40]. We obtained minor allele frequencies (MAFs) from the gnomAD database of all variants within peak regions of 31 epigenomic profiles. We observed a clear negative correlation between MAFs and MetaChrom scores, with high scoring variants present at lower MAFs (Figure 3B for two fetal and two adult cell types, the complete results in Figures S7, S8). This results thus support the deleterious effects of MetaChrom predicted functional variants.

**Fig. 1:**
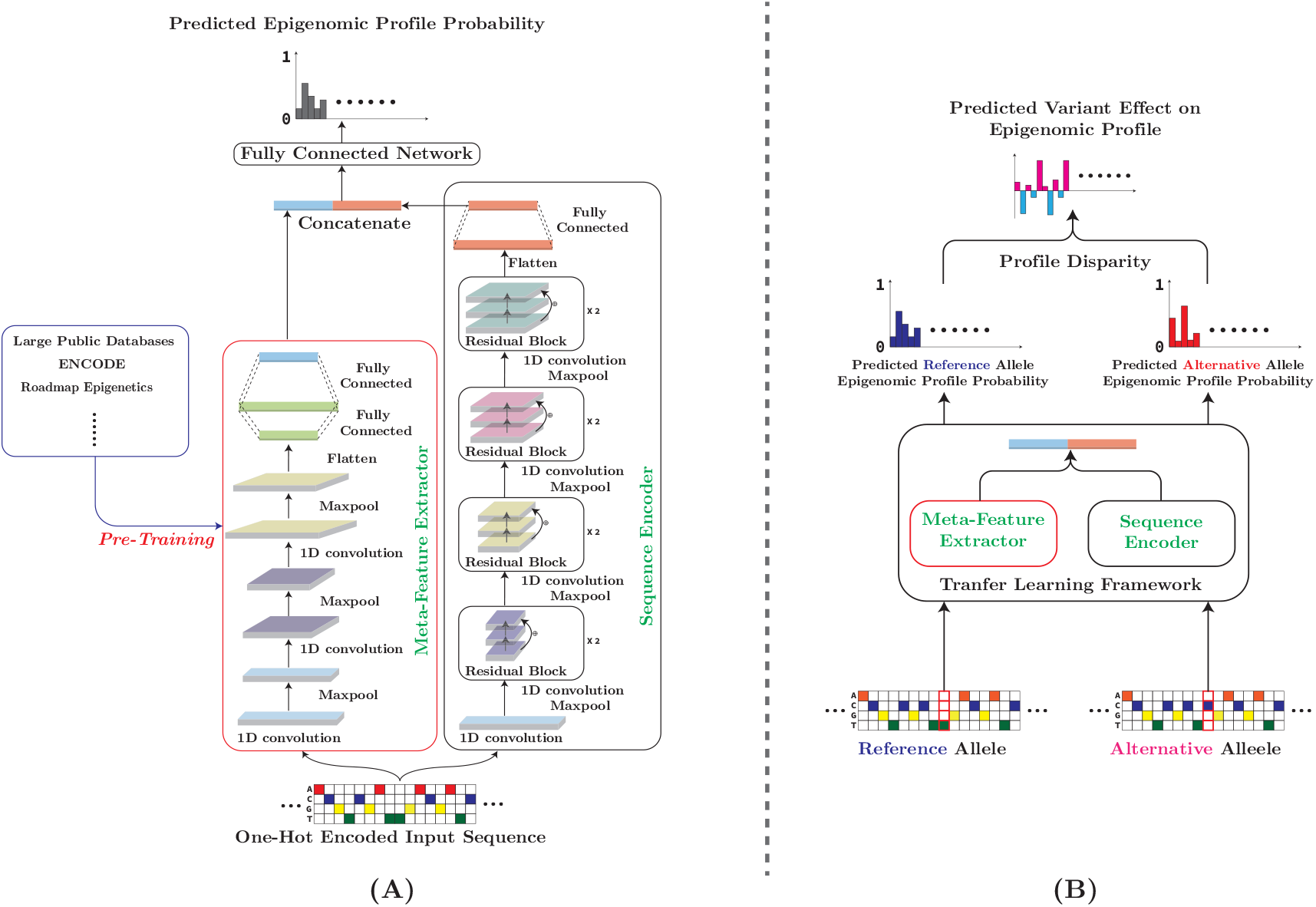
(A) Overall architecture of MetaChrom. The input sequence is fed into both MetaFeat and the ResNet sequence encoder. Their outputs are then concatenated for the prediction of epigenomic profiles.(B) Pipeline for predicting variant effect on sequence epigenomic profiles.

**Fig. 2:**
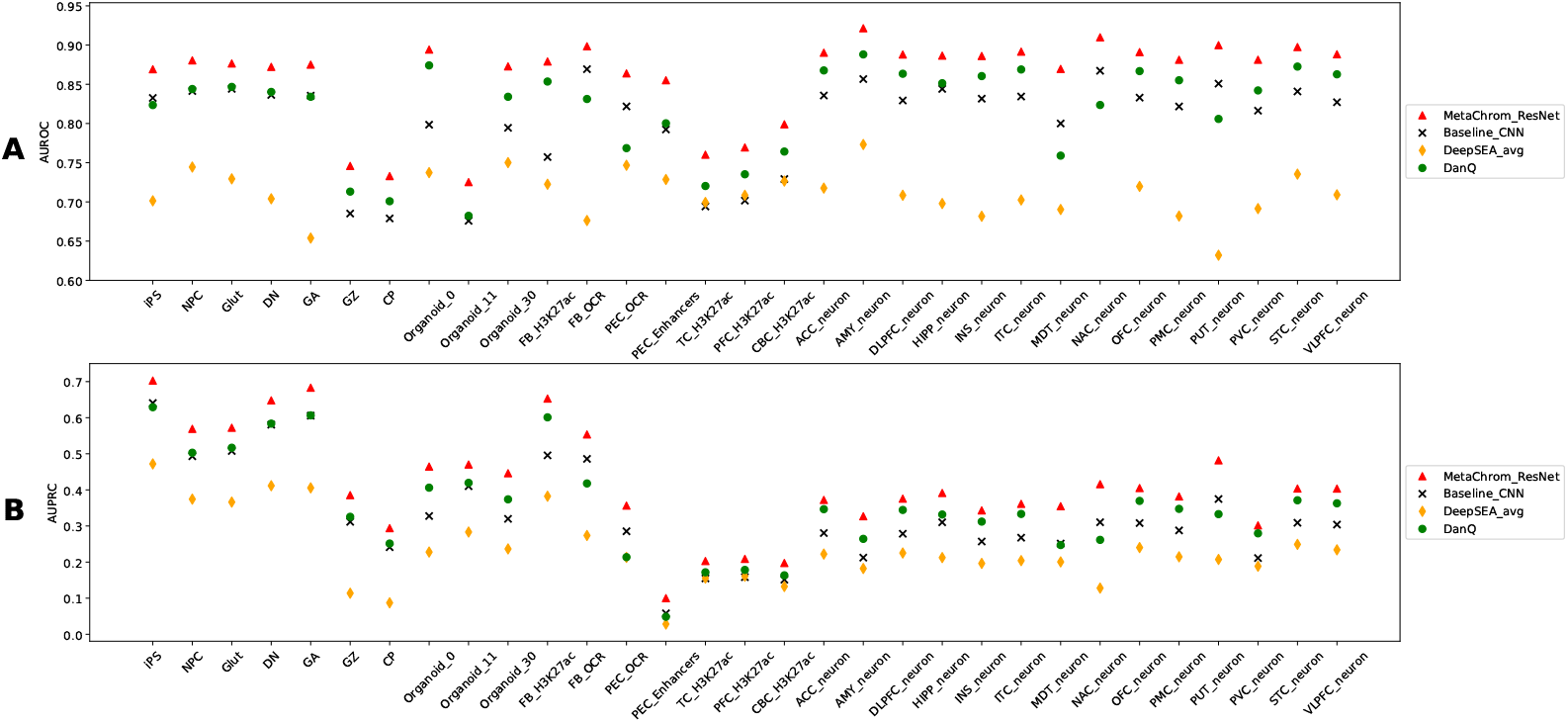
(A) AUROC and (B) AUPRC performance comparison of MetaChrom and other methods across 31 epigenomic features. See Table S1 for the list of cell/tissue types.

**Fig. 3:**
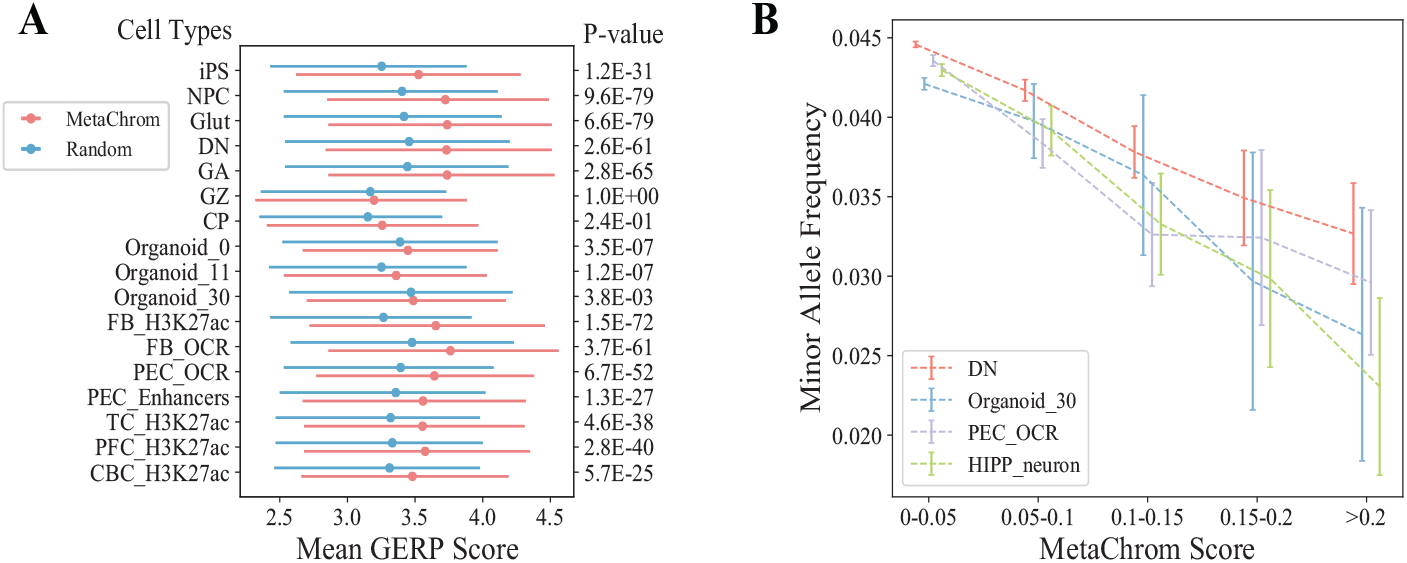
Validation of MetaChrom predicted functional variants with evolutionary constraint. (A) Distribution of GERP scores between MetaChrom predicted functional variants and random variants. (B) Minor allele frequencies of variants defined by MetaChrom scores in four selected cell types. Only variants inside peak regions of the epigenomic data were considered.

To further validate MetaChrom, we compare its predictions with experimentally determined regulatory variants in iPSC-derived neurons, based on allele-specific chromatin accessibility (ASC) analysis [90]. ASC variants are defined by allelic imbalance in ATAC-seq experiments, potentially affecting chromatin accessibility and hence gene expression. These ASC variants in iPSC-derived neurons were shown to be highly enriched with variants associated with gene expression, histone modification, DNA methylation, and risk variants of neuropsychiatric traits [90]. We focused on ASC variants from neural progenitor cells (NPC) and glutamatergic (iN-Glut) neurons, the two cell types with largest number of identified ASC variants. For all common single nucleotide variants (SNVs) in open chromatin regions of these two cell types, we computed their MetaChrom scores trained from the matched cell types. The top ranked 1,000 variants show about 6 fold enrichment of ASC variants, comparing with randomly sampled variants in open chromatin regions (Figure 4A). We also observed that MetaChrom scores from matched cell types generally show higher enrichment than other cell types, confirming the cell type specificity of MetaChrom scores (Figure S9). For comparison, we also ranked variants within open chromatin regions by CADD scores and FunSig [91, 61]. CADD score is widely used to predict deleteriousness of variants, and Funsig is a measure of predicted regulatory effects, based on deep learning models trained from a large compendium of cell/tissue types (most are not from brain). Top variants by both CADD and FunSig show modest enrichment in ASC, but at levels lower than MetaChrom (Figure 4A). These results thus highlight the power of MetaChrom in prioritizing functional variants in neurodevelopmental and neuropsychiatric contexts.

**Fig. 4:**
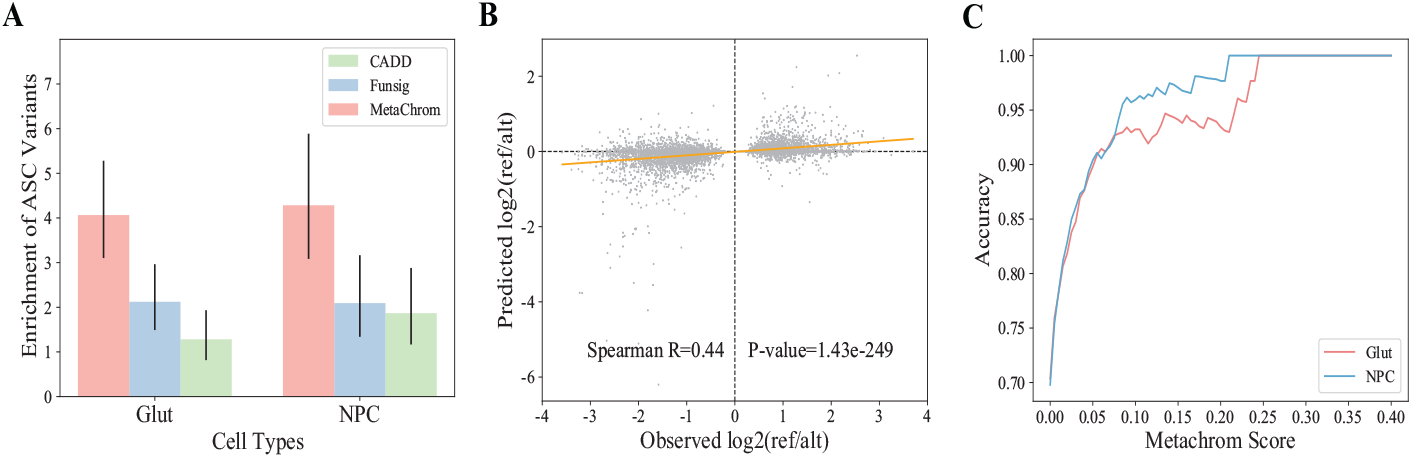
Validation of MetaChrom predicted functional variants with ASC variants. (A) Enrichment of ASC variants for predicted functional variants identified by MetaChrom, Funsig and CADD score in iN-Glut and NPC cells. (B) The observed allelic imbalance vs. MetaChrom predicted effects on chromatin accessibility of ASC variants in Glut neurons. (C) Accuracy of predicting directions of ASC variants in Glut and NPC cells.

To test if MetaChrom can accurately predict the effect sizes and directions of variants on chromatin accessibility, we compared the observed allelic imbalance of ASC variants in NPC and iN-Glut with the predicted differences between reference and alternative alleles. MetaChrom predictions track the observed allelic imbalance ratio with Spearman correlations of 0.45 and 0.40 in two cell types, respectively (Figure 4B, S8). Focusing on iN-Glut cells, we found 70% ASC variants show consistent signs in observed allelic imbalance and estimated effects (Figure 4B). ASC variants that are predicted to have large effects by MetaChrom show even higher agreement of predicted and observed directions of allelic imbalance. At MetaChrom score > 0.05, the agreement reaches nearly 90%, and goes even higher with higher MetaChrom score cutoff (Figure 4C). Together, these analyses show that MetaChrom is able to predict quantitative regulatory effects of genetic variants.

### 2.4 MetaChrom predictions of variant effects assist interpretation of GWAS results

A single locus associated with a trait from GWAS could harbor hundreds of variants in linkage disequilibrium (LD), making it difficult to distinguish causal from non-causal variants. Recent work, including our own, have demonstrated that variants disrupting chromatin states are enriched with GWAS causal signals [24, 90, 8]. Motivated by this observation, we use MetaChrom to identify putative causal variants of SCZ-associated loci. We score all the common SNPs by the absolute differences of MetaChrom predictions between two alleles in each of the 31 cell types we study. This score is then used to rank candidate SNPs from statistical fine-mapping analysis of SCZ associated loci, known as credible sets. The strength of statistical evidence of a SNP in the credible set is quantified by Posterior Inclusion Probability (PIP), Bayesian posterior probability that a SNP is a causal variant given the GWAS data. PIP values are generally small even in high-confidence GWAS loci, reflecting the uncertainty of causal variants due to LD. The PIP values are used in combination with MetaChroms scores to prioritize candidate SCZ causal variants.

We performed analysis on all SNPs in the credible sets of 145 loci associated with the risk of SCZ (Table S2). We identified 31 candidates in 30 SCZ-associated loci, based on several criteria: (i) relatively high confidence of being SCZ risk variants (PIP > 0.1), (ii) MetaChrom scores ranked at top 1% across all common SNPs in at least one cell type, and (iii) MetaChrom scores are the very top among all SNPs in the credible set of a given SCZ risk locus, in at least one cell type (Figure 5). The list includes several high confidence SNPs with PIP > 0.5. Our results thus provide further support of the disease relevance of these SNPs, and additional information about how they may function, in terms of the relevant cell types and the epigenomic features they target. The majority of SNPs have moderate PIP values (0.1 to 0.5), and would not be considered causal SNPs by themselves. Using cell type specific MetaChrom scores, we can learn the biological context through which these variants work. Based on the cell types in which a SNP’s MetaChrom score is highest among all SNPs in the credible set, we classify a SNP as acting mostly in fetal stage (F) or in adult stage (A) or both (FA). Roughly equal number of SNPs in our candidates are classified as F or A, and six SNPs as FA. These findings are consistent with a recent report that expression associated variants of fetal and adult brain make comparable contributions to SCZ heritability [80]. Interestingly, once the stage (F or A) is given, the MetaChrom scores are often not very cell type specific. Most variants acting on adult stage have high scores across multiple types of adult neurons (Figure 5).

**Fig. 5:**
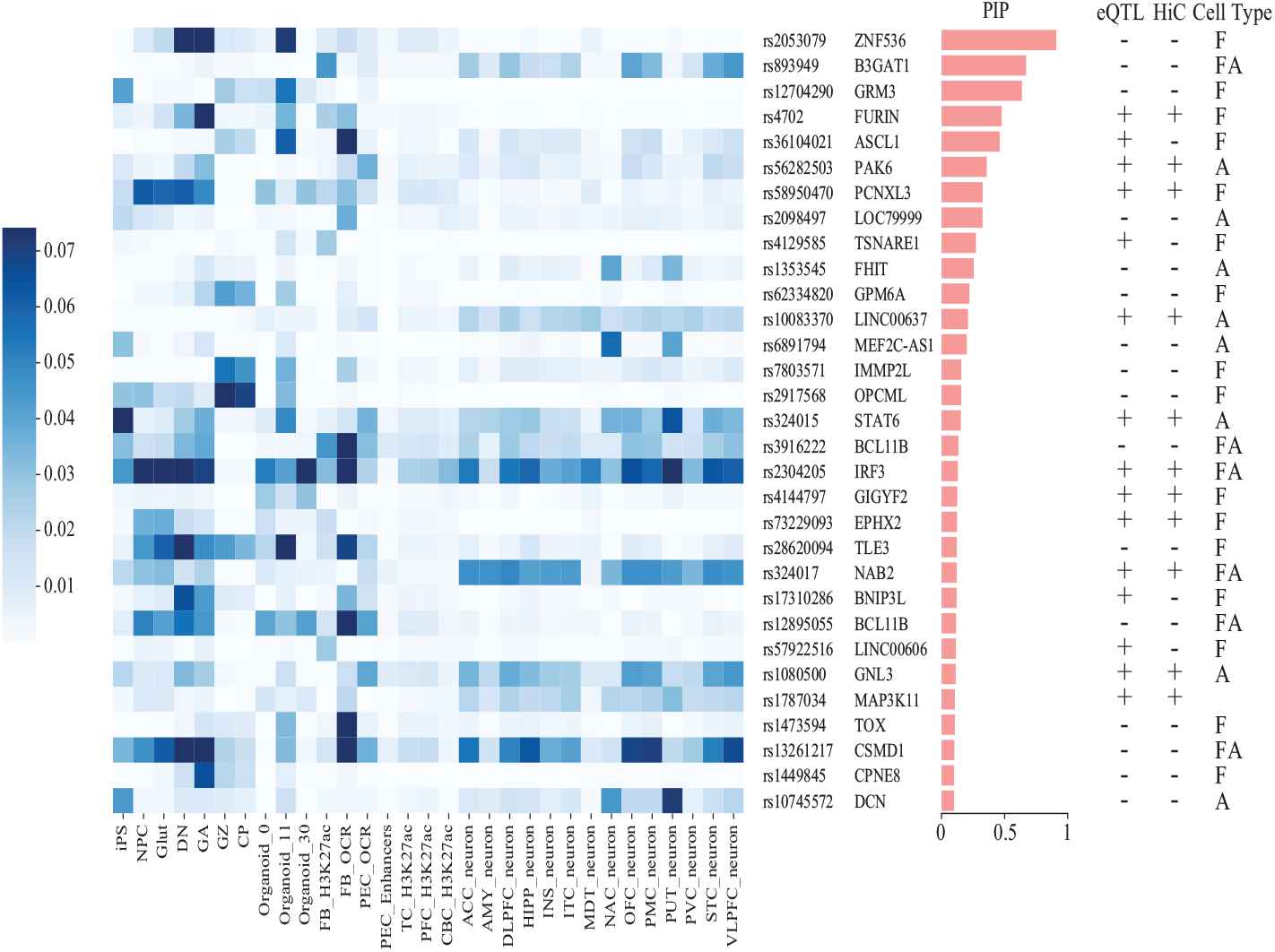
MetaChrom score of 31 candidate SNPs across 31 cell types. Candidate SNPs are ordered by their posterior inclusion probability (PIP) values shown in the middle. Three columns on the right indicate if a SNP is an eQTL (+/−), if a SNP has HiC targets (+/−) and if a SNP acts mostly in fetal stage (F) or in adult stage (A) or both (FA)

Even if causal variants are identified, their target genes may not be clear because of the possibility of long-range regulation. To assign putative target genes, we leverage external data including brain expression quantitative trait loci (eQTL) from post-morterm brain in GTEx [13] and CommonMinds Consortium (CMC) [20], and promoter-capture Hi-C data in iPSC-derived neurons [71]. A large fraction of our SNPs can be associated with one or more genes in eQTL or Hi-C data (Figure 5, Table S3). Additionally, a few SNPs are located in the promoter and UTR regions of genes. Combining these evidences and literature search, we assign the most likely target genes at each of the 31 SNPs (Figure 5, Table S3). A number of these genes represent highly plausible risk genes of SCZ. For instance, FURIN and TSNARE1 were recently confirmed to regulate neuron growth and synaptic development by CRISPR editing in iPSC derived neurons [64]. GRM3 is a glutamate receptor and is being explored as a therapeutic target of SCZ [23]. ZNF356 is a transcription factor with an essential role in development of a small subset of forebrain neurons implicated in stress and social behavior [75].

We discuss the SNP, rs2304205, in depth to show how our method may assist the study of genetics of complex traits (Figure 6). The SNP is located in a region strongly associated with SCZ, with multiple SNPs having *p*-values below genomewide threshold (Figure 6, top). Statistical fine-mapping is insufficient to resolve the causal variant, as the credible set contains 12 SNPs with PIPs of all below 0.2. MetaChrom analysis highlights rs2304205 as the most plausible causal variant in the credible set. It has high scores across almost all cell types, in both fetal and adult stages (Figure 5). In 24/31 cell types we examined, rs2304205 has highest MetaChrom scores among the SNPs in the credible set. The scores in four of these cell types, two each in fetal and adult stages, were shown in Figure 6. The SNP is located in the UTR regions of both IRF3 and BCL2L12. Only IRF3 is found as the likely target gene of rs2304205 in brain eQTL data (Table S3). IRF3 is a key regulator of the innate immune system and is a plausible SCZ risk gene [88]. Recent studies show that it may be important in regulating the development of neuronal progenitor cells [48], and physically interacts with other schizophrenia susceptibility genes, such as CREB1, AKT1 and ESR1 [88]. We performed additional in-depth study of a region containing rs1080500, which is highlighted as possible causal variant by MetaChrom. The SNP effect is likely limited to adult neurons and is also an eQTL in adult brain (Figure S11). The likely target gene based on eQTL, GNL3, is known to regulate neuron differentiation [55]. Taken together, these case studies highlight the potential of MetaChrom in prioritization of causal variants following GWAS.

**Fig. 6:**
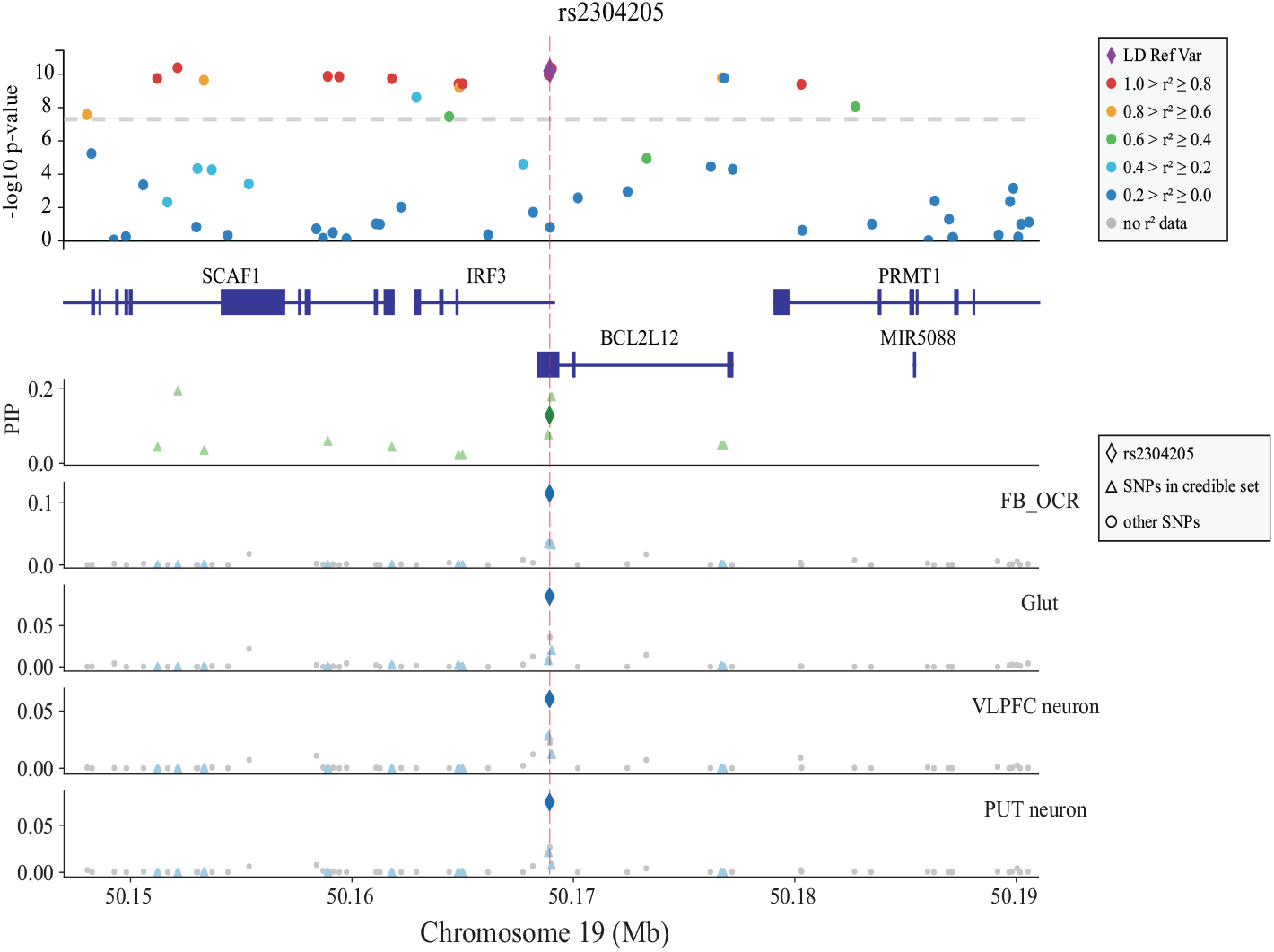
Likely causal variant rs2304205 and its MetaChrom functional annotations. The candidate SNP rs2304205 is chose as the reference variant for computing LD and it is highlighted by the red dash line in each panel. The upper panel shows significance of GWAS SNPs, LD between SNPs and genes in this region. The next panel shows credible set SNPs identified by fine-mapping (PIPs) in this region. The remaining panels show MetaChrom scores in four cell types, two in fetal stage (FB OCR and Glut) and two in adult stage (VLPEC neuron and PUT neuron)

### 2.5 Identification of biologically relevant motifs from MetaChrom

To better understand what have been learned by our model, we extracted representative sequence patterns from our model using TF-MoDISco[67] and TOMTOM[29] (See Methods). We first sampled DNA sequence bins from our test set with mutually exclusive epigenomic feature labels. That is, each sampled sequence fragment is marked positive in only one of the 31 epigenomic features. Then we computed the gradient with respect to the input sequence for cell-type-specific saliency signals and used it as the input to TF-MoDISco. The sequence patterns generated by TF-MoDISco are then matched to known TF (transcription factor) binding motifs at human CIS-BP[85] using TOMTOM[29].

As shown in Fig. 7, our model detected known TF binding motifs specific to certain epigenomic features as well as motifs shared by multiple epigenomic features. For example, we detected binding motifs of the FOS and JUN families that form the activator protein 1 (AP-1) complex across various epigenomic features. These protein and protein complex are known to be associated with brain development[89, 79]. We detected many iPSC-specific motifs such as CTCF, an important regulator for chromatin structure[53], and POU5F1B(OCT4-PG1), a key player in the stem cell induction process[73]. In dopaminergic (DN) cell, we found motifs of proneural transcription factor FOXP4 that plays an important role in neural development[44] and FOXO6, a transcription factor closely related to cortical development[54, 72]. We discovered OLIG2, a transcription factor associated with cortical neurogenesis[45] and EN2 a transcription factor links to many stages of neural devlopment[25] in samples from the human neocortex(GZ, CP)[16].

**Fig. 7:**
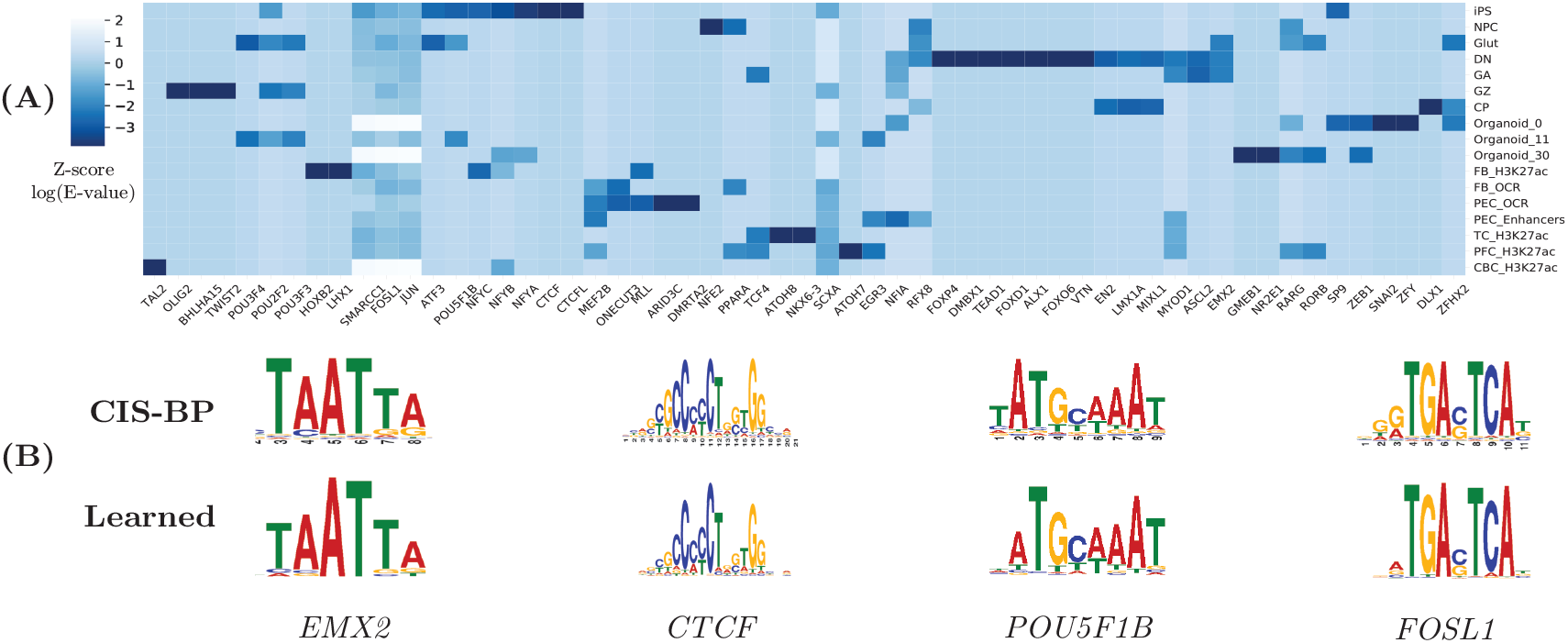
(A)Distribution of matched motifs in each epigenomic assay and (B)selected binding motifs identified by our method and their matches in the CIS-BP database

## 3 Discussion

In this paper we have presented a novel deep transfer learning method, MetaChrom, that predicts epigenomic features from only DNA sequences, by leveraging large publicly available databases of epigenomic profiles. By coupling ResNet[30] with transfer learning, we have significantly improved over previous CNN models the prediction accuracy of epigenomic profile in a set of neurodevelopmental epigenomic profiles. Using a combination of evolutionary constraint and experimentally determined ASC variants, we validated the utility of MetaChrom in predicting functional effects of variants, and illustrated how MetaChrom facilitates the prioritization of causal risk variants in GWAS loci associated with SCZ. We note that while the model was developed in the context of neurodevelopment, the approach and software is generic and can be applied to any user-provided epigenomic data in other biological contexts.

We note an important limitation of our and related machine learning methods for predicting functional effects of variants. The objective of these methods is to predict epigenomic features, e.g. chromatin openness of a region, which is a different task from predicting variant effects. Regulatory sequences may differ from background sequences in systematical ways. Studies have shown that mutational processes in DNA sequences depend on nucleosome positioning and binding of proteins [42, 9, 62], which are generally different between enhancers and other sequences. Over evolutionary time, these different mutational processes may lead to accumulation of distinct sequence features in regulatory sequences, which would not be useful for predicting functions of variants. Another problem of training the models using epigenomic profiles is that regulatory sequences may be enriched with motifs that are important in other biological contexts, e.g. when cells are exposed to certain simulation, or during early developmental stages[81]. Whether these contexts are relevant to the phenotypes of interest may be unclear. As a result, not all sequence features learned by the machine learning methods would be useful for predicting functional variants in the current context/conditions. We think an important direction is to make better use of experimentally determined and context-specific, functional variants during training of machine learning models.

To evaluate variant scoring tools, current work usually relies on two sources of known functional variants: known pathogenic variants such as those from HGMD [91] and expression QTLs (eQTLs). Each has its own limitations. The number of known pathogenic variants in a given phenotypic context is often very limited. For eQTLs, they are often variants in LD with true functional variants, but not functional ones themselves. In our work, we overcome these limitations by taking advantage of a large collection of ASC variants in iPSC-derived neuronal cells. Unlike eQTLs, ASC variants are inside or very close to open chromatin regions, and most of them are expected to be functional [90]. Using this dataset, we demonstrated that our MetaChrom predicted variants are much more likely to be ASC variants than methods that do not take into account specific biological contexts (Figure 4). Interestingly, we note that despite our superior performance, only about 20-27% of MetaChrom predicted variants are ASC variants (Figure S9, see Methods). This may partly reflect the limited power of the experimental study, as many more ASC variants may be missed. But it also highlights the challenge of predicting functional variants, as opposed to predicting epigenomic profiles. In fact, the method performs very well in classifying open chromatin regions, with AUROC and AUPRC close to 90% and 60%, respectively (Figure 2), but the accuracy of predicting ASC variants is still rather modest. Together, our efforts using ASC highlight the benefit, as well as challenge, of evaluating computational models using experimentally determined variants.

In analysis of GWAS data of SCZ, we demonstrated that MetaChrom predictions help prioritize causal risk variants and by combining with other datasets, reveal mechanistic insights of these variants. One limitation is that we do not yet have a single quantitative metric that combines statistical associations with deep learning based functional predictions. In our recent study [90], we show that it is possible to use ASC variants as functional information as prior in Bayesian fine-mapping. It would be interesting to extend such strategy to MetaChrom predictions. One possibility, for instance, is to combine MetaChrom predictions with experimental ATAC-seq data to have better power of detecting ASC variants. Such functionally informed genetic variant mapping has been used in eQTL studies and GWAS [86, 57]. The resulting set of deep learning-enhanced ASC set, when used as prior, may provide even better resolution for fine-mapping GWAS loci.

In conclusion, we developed a deep learning based tool for predicting functional genetic variants in neurodevelopmental context. We demonstrated its accuracy using known regulatory variants in neuronal cells and its potential of revealing risk variants of GWAS of mental disorders. This tool is generally applicable and may enable researchers to better translate GWAS associations into mechanistic insights.

## 4 Methods

### 4.1 Reference epigenomic profile data

We downloaded the epigenomic profile dataset from the DeepSEA website (http://deepsea.princeton.edu) [91]. This dataset consists of 919 chromatin features derived from the ENCODE and Roadmap Epigenomics [12, 5]. The epigenomic features are computed by first binning the reference genome (GRCh38/hg38) into 200-bp sequence fragments. The fragments were then intersected with the downloaded peaks from the public databases. Each bin was assigned a binary vector *l* ∈ *R*^*d*^(*d* = 919) as its label, each dimension representing an epigenomic feature *i* from a specific cell type with corresponding sequencing assay. If a fragment is at least 50% overlapped with the peak present in the sequencing assay *i*, the corresponding dimension in *l* is assigned 1 (i.e., *l*_*i*_ = 1), otherwise 0 (i.e., *l*_*i*_ = 0). After computing the features, the fragments were extended to 1kb to include surrounding sequences [91, 3]. All the fragments were then split into training, validation, and test sets such that all fragments from chromosomes 7 and 8 are held out for test, and the rest are randomly split into training and validation sets. The training set we used contains 4,400,000 sequence fragments with at least one positive chromatin feature. We have trained our meta-feature extractor (MetaFeat) on this chromosomal-based split so we can fine-tune the feature extractor using the test sequence. We also employed the same chromosomal-based split in our case study for neurodevelopment related tissues data(Method, Table S1) which ensures the test sequences are never seen by our model before test time to avoid potential training bias.

### 4.2 Epigenomic profiles from neurodevelopment related tissues and cell types

To comprehensively capture the epigenomics landscape of the human neurons and brain, we collected data from 31 different epigenomic assays, from the early developmental stages to fully developed adult brain tissues. For the early developmental stages, we obtained a set of ATAC-seq peaks from iPSC derived neuronal cells described in[90], with a total of five different cell types. These cell types are good models of neurodevelopment. We also obtained one fetal brain DNase-seq sample from the Roadmap Epigenomics Project [37]; chromatin accessibility data from brain organoid samples at three different time points [76]; ATAC-seq profiles from two early human neocortex samples in germinal zone (GZ) and cortical plate (CP) [17]; and one fetal brain H3K27ac profile from [60]. For the adult brain, we collected fourteen neuronal ATAC-seq profiles from the BOCA project [22]; and five chromatin and histone features from the PsychENCODE project [82].

We processed the peaks from each epigenomic profile in a similar way as the aforementioned reference epigenomic profile data. In our dataset, each dimension in the label *l* ∈ *R*^31^ represents if a segment is active or not in a given epigenomic profile, i.e., if it is at least 50% overlapped with the peaks in each epigenomic assay. We choose our test set such that all fragments from chromosomes 7 and 8 are held-out when the rest of the genome are randomly split for training and validation. After processing, we obtained 3,165,290 sequence fragments that’s active in at least one epigenomic assay for training and validation; our test set contains 390,380 sequence fragments for model evaluation.

### 4.3 MetaChrom model architecture

MetaChrom as shown in Fig. 1(A) has two major modules: 1) a CNN-based meta-feature extractor pre-trained on a large public dataset(MetaFeat). 2) a ResNet-based sequence model that learns cell-type-specific features directly from the input sequence. To predict the regulatory profile of a DNA sequence fragment *i* of 1kbp in length, we first encode the sequence fragment as a one-hot matrix and fed it into the meta-feature extractor, which will output a vector representation 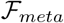 of the input sequence while the sequence is also fed to the ResNet-based sequence encoder simultaneously to obtain a cell-type-specific feature representation 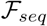. The two feature representations are concatenated to form a new joint vector representation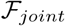, which is fed into a fully connected dense network to predict epigenomic features 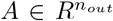, Where *n*_*out*_ is the number of epigenomic features of interest.

MetaFeat is a CNN model consisting of three 1-D convolutional layers with kernel length 7 and channel sizes 320, 480 and 960, respectively. This feature extractor takes a one-hot encoded sequence (of dimension 1000 × 4) as input and outputs a vector representation(*R*^919^) of the input sequence fragment. Each convolutional layer in this feature extractor is followed by a ReLU [50] layer for activation and a max-pooling layer with a kernel size of 4 for down-sampling. We used a smaller kernel size (7), while previous methods use a large kernel size (19) for model interpretability [32], but it has been shown that with appropriate interpretation tools [67, 49] meaningful binding motifs may be detected with a smaller kernel. The final convolutional layer connects to two fully connected layers, which in turn generate a vector of 919 chromatin features to represent the input.

Our ResNet-based sequence encoder maps the input sequence (encoded as a one-hot matrix of dimension 1000 × 4) to a cell-type-specific representation, which is a vector of 31 elements. The ResNet model consists of one 1D convolutional layer and eight residual blocks. The 1D convolutional layer has a kernel length 4 and channel size 48. Each residual block contains two 1D convolutional layers, each followed by a ReLU activation layer. The eight residual blocks have channel sizes of 96, 96 128, 128 256, 256, 512 and 512, respectively and kernel length 7. Finally, the last residual block connects to two fully connected layers, which generate the cell-type-specific representation for the input sequence segment.

The outputs from MetaFeat and the sequence encoder are concatenated to form a integrative vector representation of the input sequence, which is then fed into three fully connected layers to predict the probabilities of epigenomic profiles.

### 4.4 Predicting Variant effects on regulatory profiles

To predict the variant effect on a given sequence’s epigenomic profile, two sequences of length 1kb differing only at and centered at the variant position are used. One of them corresponds to the reference allele and the other corresponds to the alternative allele. As shown in Fig. 1(B), we pass those two sequences into MetaChrom separately to predict their epigenomic profiles as *A*_*ref*_ and *A*_*alt*_. Then we compare the predicted regulator profile and compute the disparity as absolute difference |*A*_*ref*_ − *A*_*alt*_| or log odds ratio 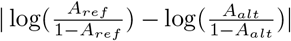 for measuring variant effects.

### 4.5 Model training and testing

We trained our transfer learning framework in two phases. In phase one we train the metafeature extractor on the public epigenomic profile data (see Data section) using binary cross-entropy as the loss function and the Adam optimizer [33]. In phase two we jointly train the ResNet model and the meta-feature extractor [43] with stochastic gradient descent(SGD) accelerated by Nesterov momentum [51], starting from the trained feature extractor in the first phase. In both phases, sequence bins from chromosome 7 and 8 are held-out for test and the training/validation sets are randomly split from the remaining bins. For model selection, we searched hyperparameters for different batch size, learning rate and channel size in phase one. In phase two, we searched for different batch size, momentum, learning and weight decay rates. All models are trained on an NVIDIA 2080Ti GPU in 5 hours.

### 4.6 Assessing evolutionary constraint on MetaChrom predicted functional variants

A variant is scored by MetaChrom using the absolute value of the difference in MetaChrom output between reference and alternative alleles. The score ranges from 0 to 1. For top MetaChrom predicted variants in 31 cell types, we calculated and compared their GERP scores [15] with control variants, chosen randomly in peak regions of the same cell types. GERP scores were obtained from ANNOVAR [83], and were compared between MetaChrom predicted and control variants using the Wilcoxon Rank-Sum Test.

Minor allele frequencies (MAFs) of all variants were obtained from the Genome Aggregation Database (gnomAD) [31] using ANNOVAR. We split variants into two sets of 5 bins based on the MetaChrom score: 0-0.05, 0.05-0.1, 0.1-0.15, 0.15-0.2, 0.2-1.0 and 0-0.05, 0.01-0.02, 0.02-0.03, 0.03-0.04, 0.04-1.0. In each bin, mean MAFs were calculated to investigate the correlation between MAFs and MetaChrom scores.

### 4.7 Evaluation of MetaChrom prediction using ASC variants

We used a recently published dataset of allele-specific chromatin accessibility (ASC) variants to compare several methods for predicting functional variants [90]. ASC variants are defined by allele imbalance in read counts from ATAC-seq experiments. A total of 5,611 and 3,547 ASC SNPs, at FDR < 0.05, were identified in neural progenitor cells (NPC) and gutamatergic neurons (iN-Glut), respectively.

To identify putative functional variants, we limit to single nucleotide variants (SNVs) in open chromatin regions in 2 cell types. The SNVs are retrieved from the 1000 Genomes Project with MAF > 5% [11]. We calculated MetaChrom score, Funsig score and CADD score for all these SNVs. Funsig scores were obtained from the DeepSEA Server [91] and CADD score [61] from ANNOVAR. We chose the top 10,000 variants ranked by MetaChrom, Funsig and CADD in descending order as predicted functional variants for each method. We then counted the number of ASC variants in predicted functional variants vs. control variants. The enrichment of ASC variants is then calculated by Fisher Exact Test.

In the analysis involving effect size and direction of SNPs, we define the observed allelic imbalance as log(*R*_*ref*_ /*R*_*alt*_), where *R*_*ref*_ and *R*_*alt*_ denote the number of reads mapped to the two alleles, and the MetaChrom predicted effects on chromatin accessibility from our model as log(*A*_*ref*_ /*A*_*alt*_). Correlation between observed allelic imbalance and MetaChrom predicted effects on chromatin accessibility is calculated by Spearman’s rank correlation coefficient.

We also estimated the percent of MetaChrom predicted variants are actually experimentally determined ASC variants. For iN-Glut cells, among top 1000 MetaChrom variants, 462 SNPs are evaluated for allele imbalance test (not all SNPs are heterozygous in the study), and 123 (27%) are reported as ASC variants. For NPC, 445 SNPs, among top 1000, are evaluated for allele imbalance test and 87 (20%) are ASC variants.

### 4.8 Motif identification and visualization

Our model learns context-specific sequence patterns to predict epigenomic profiles. Many of these patterns may correspond to cell-type-specific transcription factor (TF) binding motifs. One way to interpret the models is to extract sequence patterns (motifs) from the filters of the first convolutional layer[32, 1, 59], but this strategy often yields patterns that are hard to interpret. This is because CNNs learn a distributed representation of sequence motifs and thus, an individual filter may correspond to only a partial motif that cannot be easily identified [35, 67]. Some methods assess the importance of each position in a given sequence to measure individual nucleotide contribution[66], but they do not yield interpretable motifs directly.

To extract meaningful sequence motifs from our deep model, here we used the recently-developed tool TF-MoDISco [67] that combines position-wise single-nucleotide contribution scores to generate cell-type-specific sequence patterns. We then use TOMTOM[29] to match the identified sequence patterns to known TF binding motifs in the CIS-BP database for further analysis[85]. To apply TF-MoDISco, we first randomly select 2,000 DNA bins from our testing set with mutually exclusive epigenomic features, in which each sequence bin is only marked active in one epigenomic feature but not others. Then we generate position-wise contribution score with saliency map[68], which is the gradient of the output with respect to the one-hot encoded input. The gradient is then gated by the observed nucleotide to generate importance score, i.e., only the gradient of the observed nucleotide is kept and the gradient of unobserved nucleotide is set to 0. The importance score is fed into TF-MoDISco to generate predictive sequence patterns, which were searched against the human CIS-BP [85] database for TF binding motifs using TOMTOM [29] with E-value=1e-4.

---

**Fig. S1.**
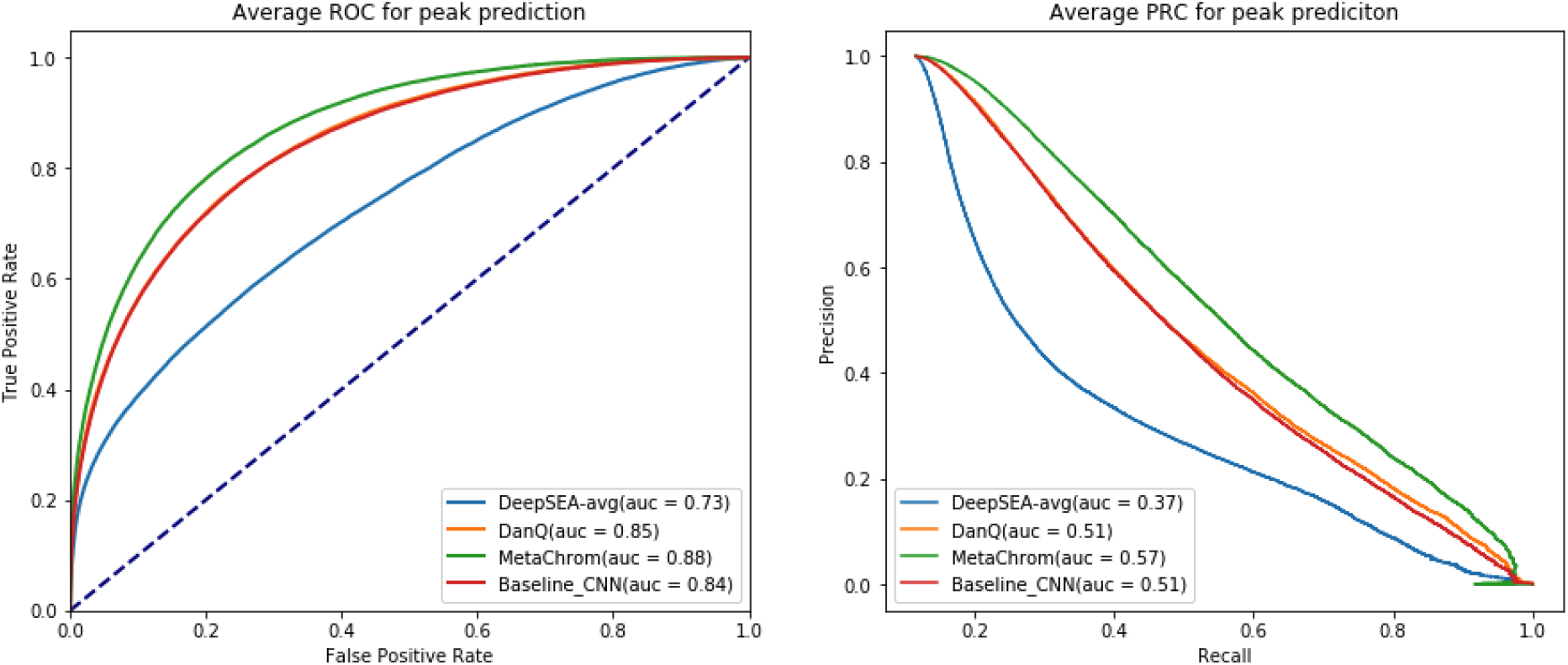
ROC and PRC plot for iPS cell derived Glutamernergic neurons acriss different tested models.

**Fig. S2.**
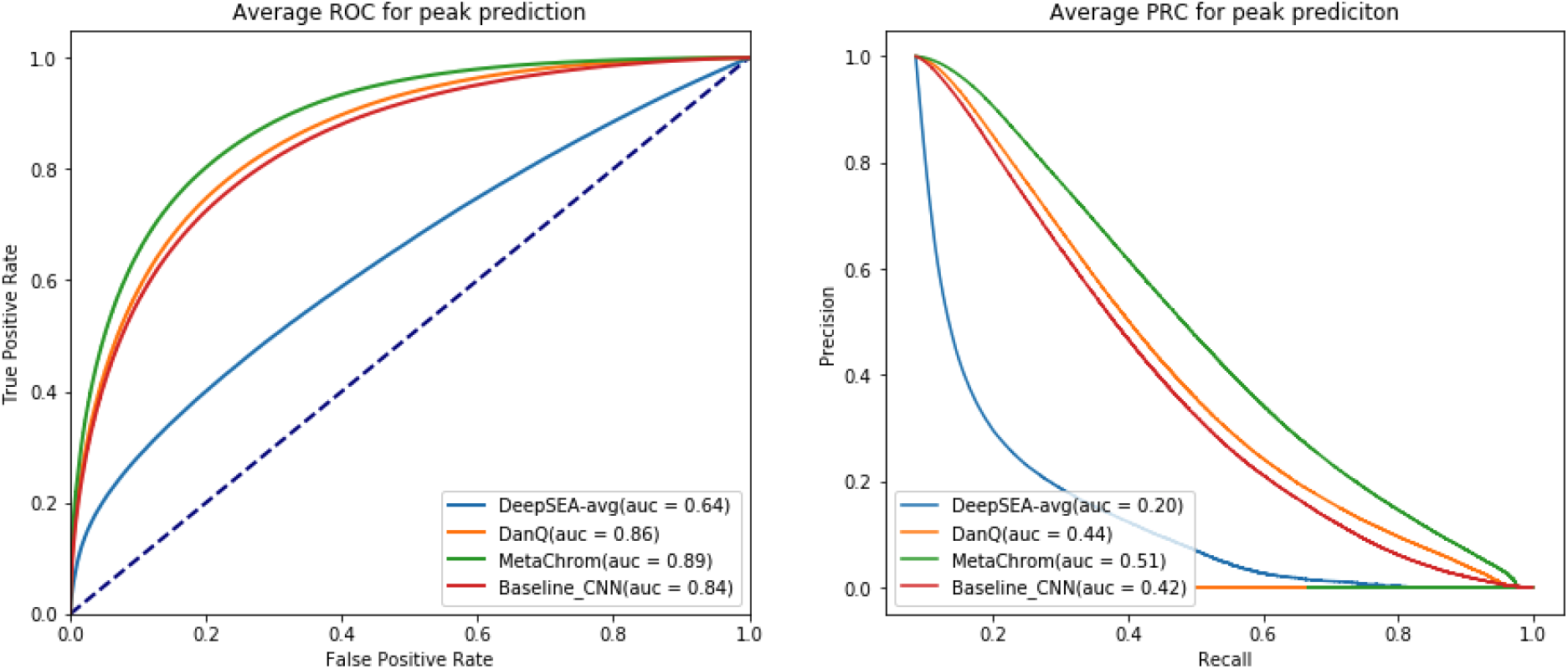
Average ROC and PRC plot for 31 epigenomic features across different tested models.

**Fig. S3:**
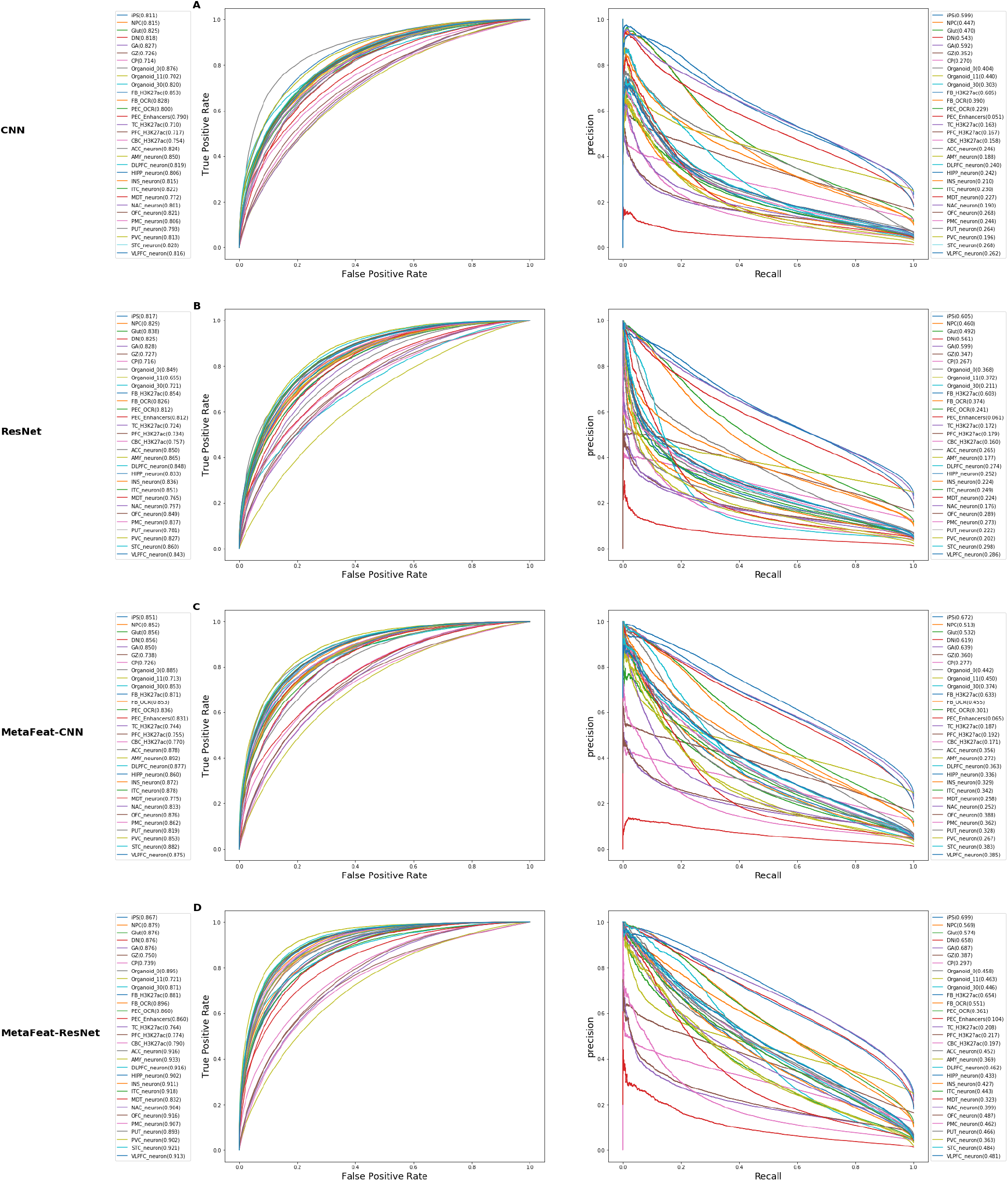
ROC and PRC plot for (A) CNN, (B) ResNet, (C) MetaFeat-CNN, (D) MetaFeat-ResNet models on 31 epigenomic features(Method).

**Fig. S4:**
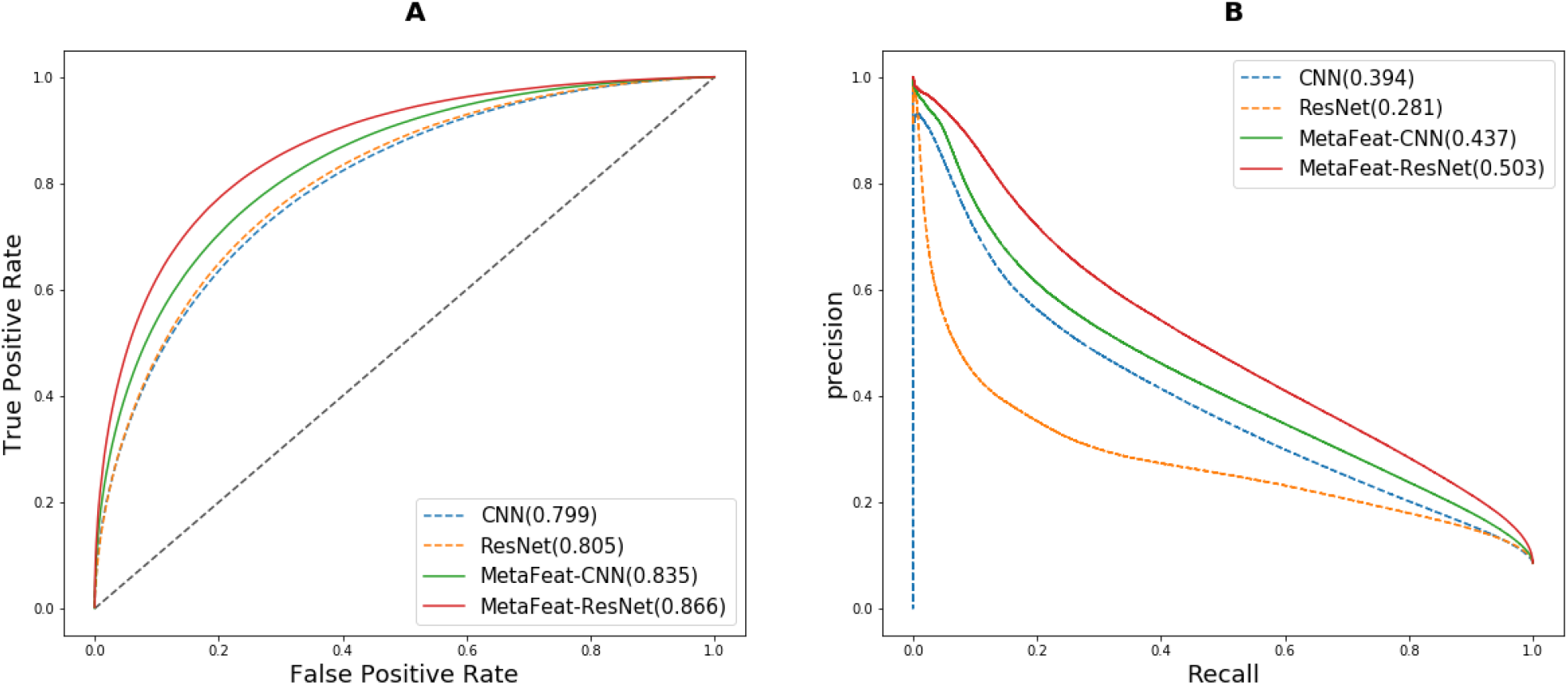
Average ROC and PRC plot for 31 epigenomic features across different tested models.

**Fig. S5:**
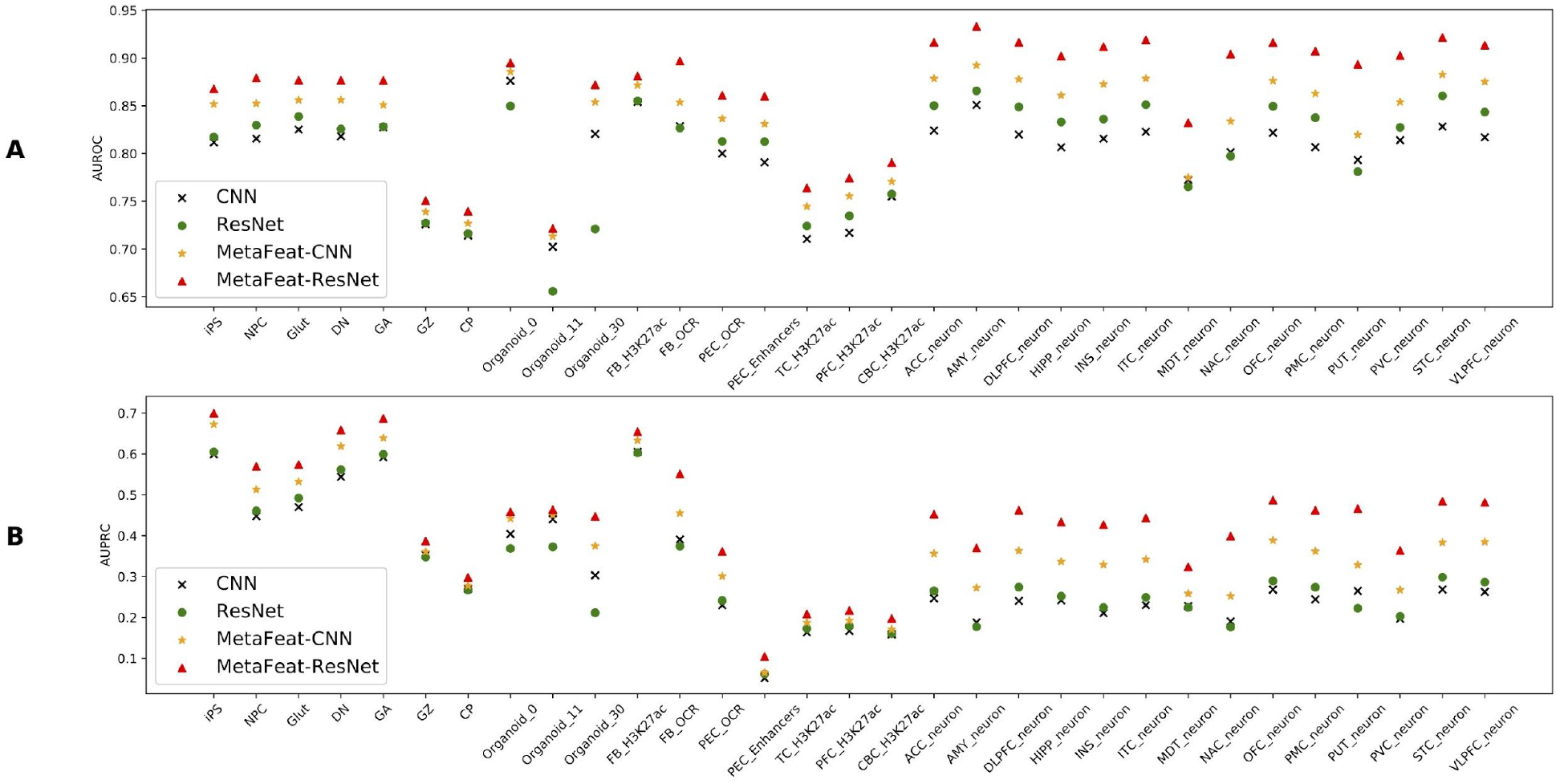
(A) AUROC and (B) AUPRC performance comparison of MetaChrom and other methods across 31 epigenomic features. See Table S1 for the list of cell/tissue types.

**Fig. S6:**
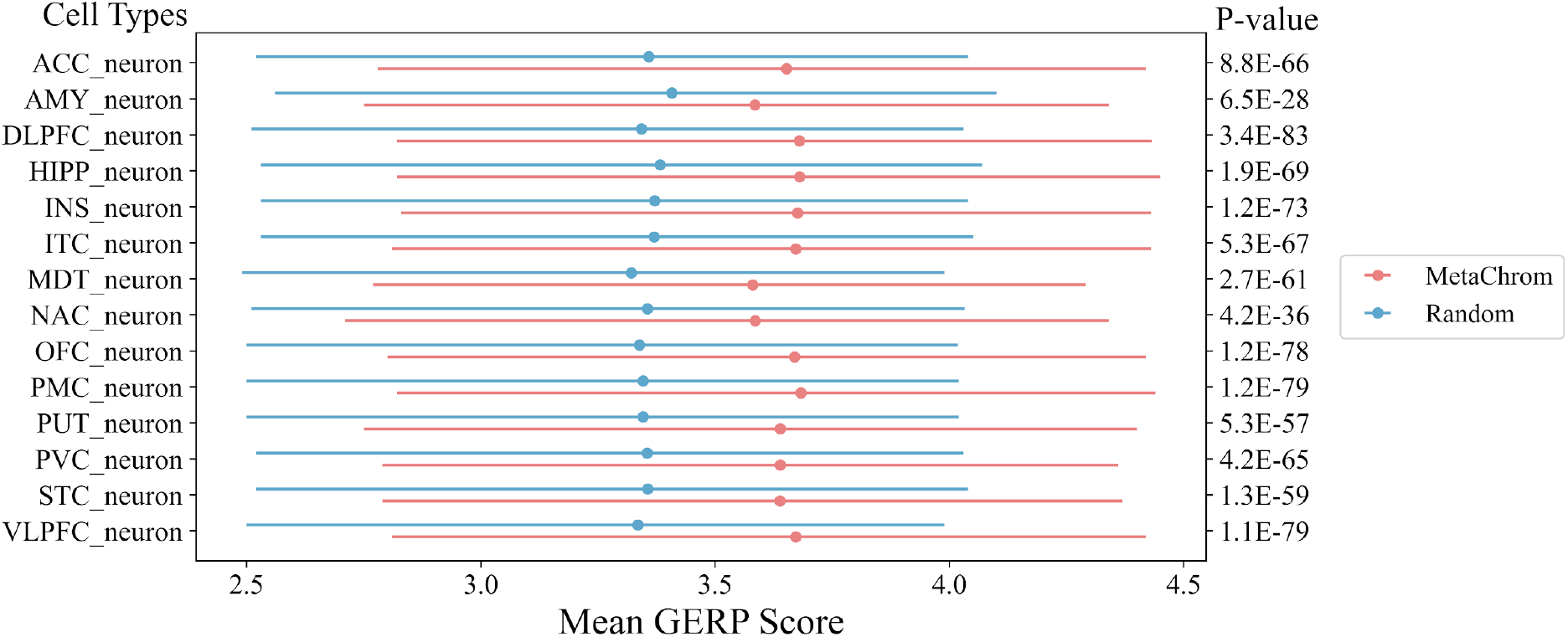
Distribution of GERP scores between MetaChrom predicted functional variants and random variants sampled from the peak regions in each cell type. P-values testing the difference were computed from Wilcoxon Rank-Sum Test.

**Fig. S7:**
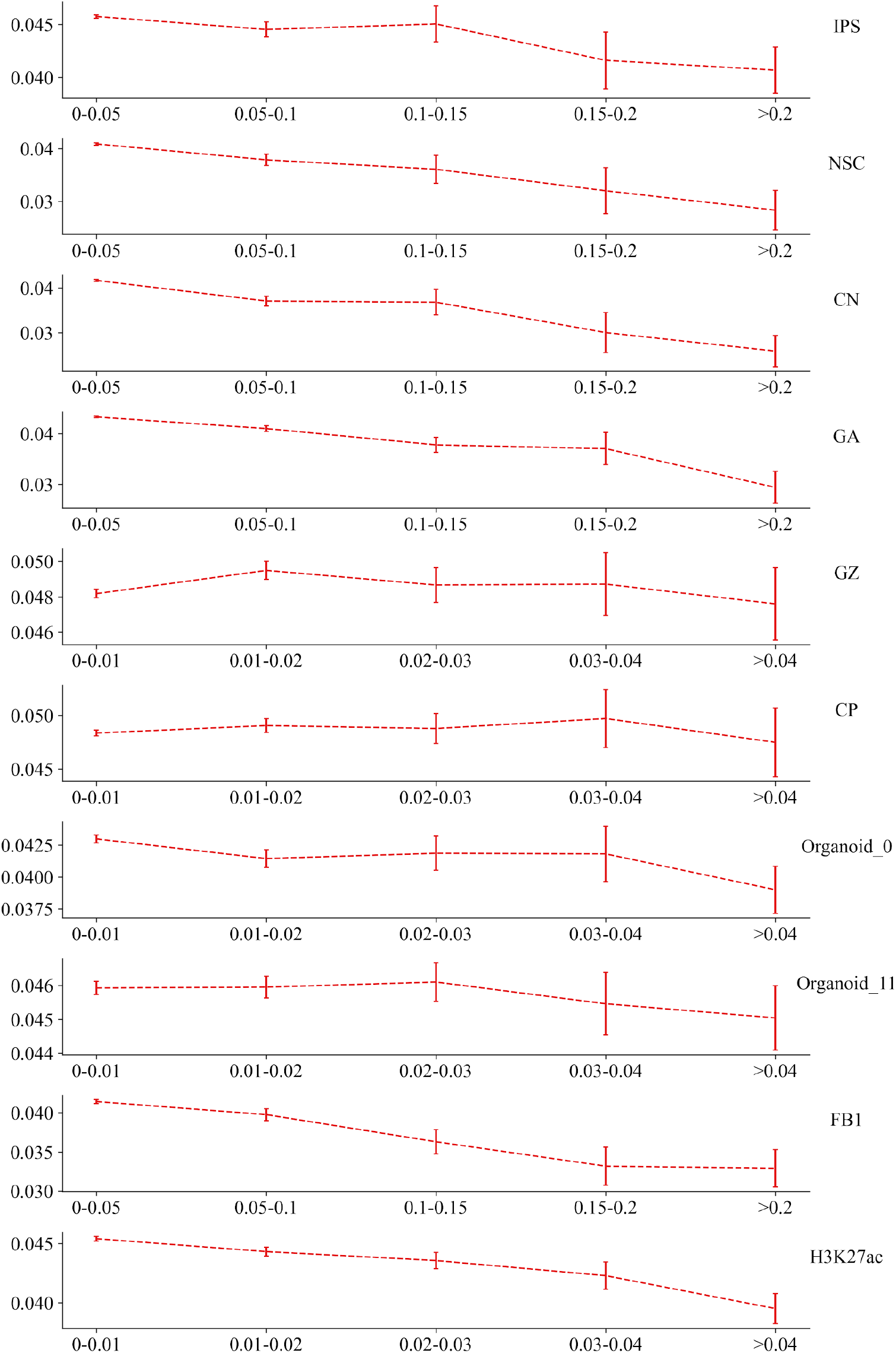
Minor allele frequencies of variants defined by MetaChrom scores in 10 epigenomic profiles in fetal brain cell types. Only variants within peak regions of the data were considered.

**Fig. S8:**
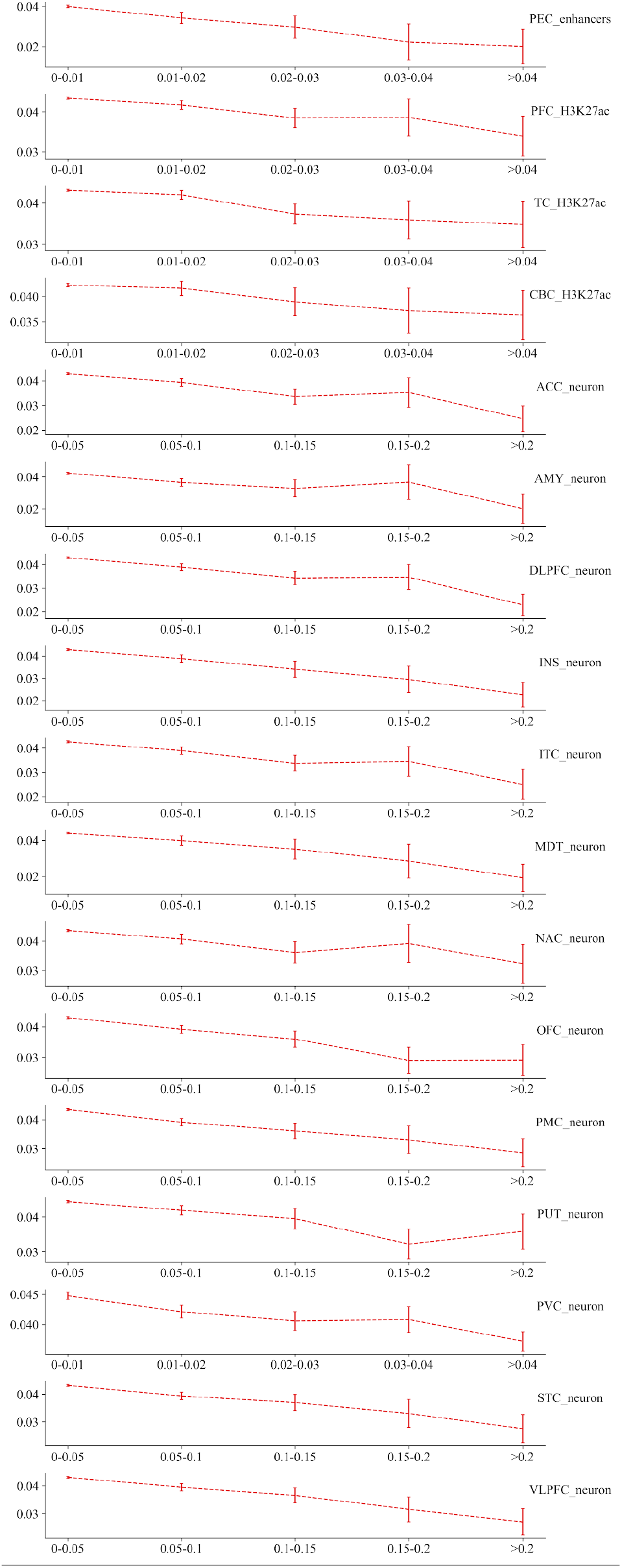
Minor allele frequencies of variants defined by MetaChrom scores in 17 epigenomic profiles in adult brain cell types. Only variants within peak regions of the data were considered.

**Fig. S9:**
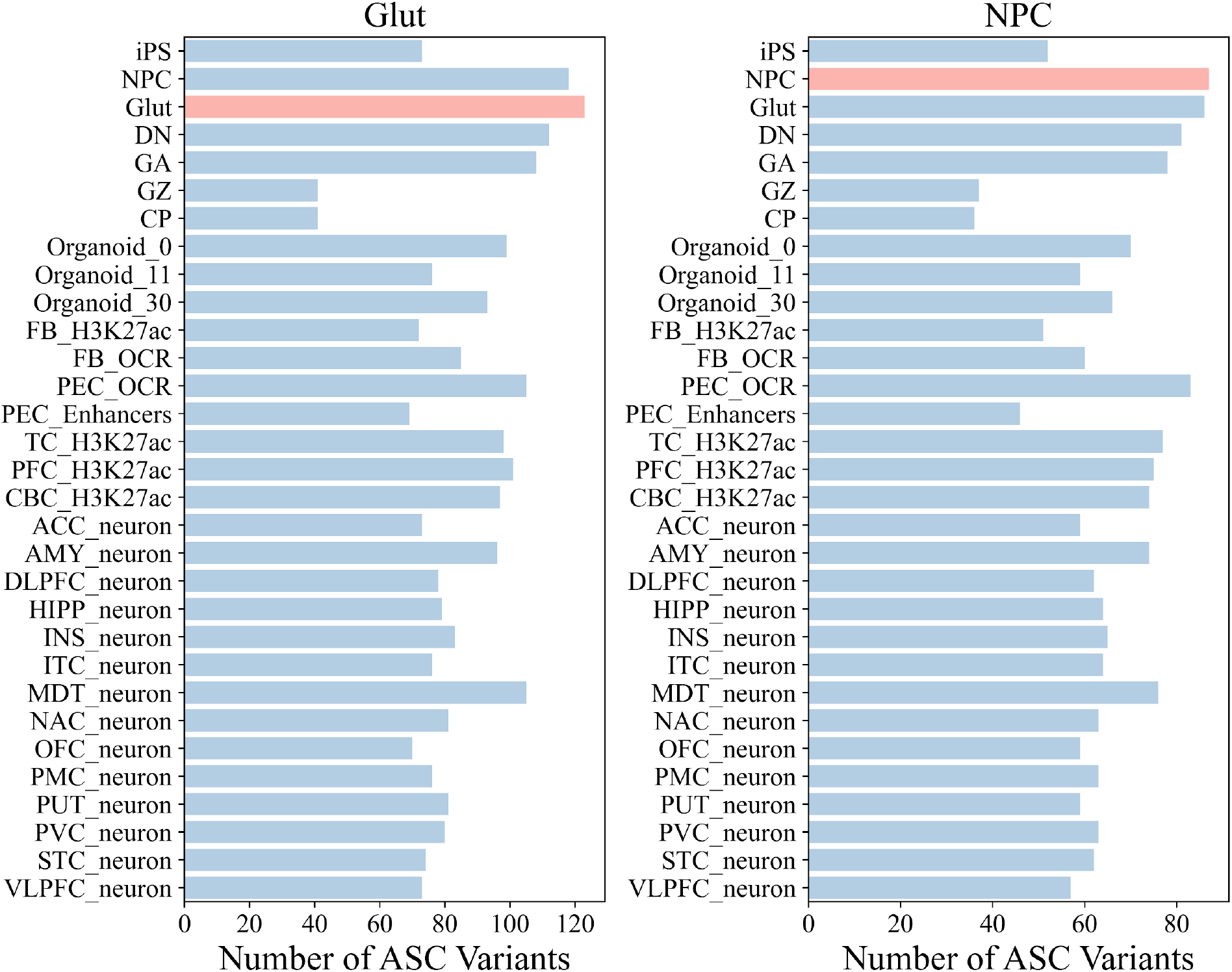
Number of experimentally determined ASC variants (two cell types, Glut - left and NPC - right) in top 10,000 MetaChrom predicted functional variants across 31 cell types.

**Fig. S10:**
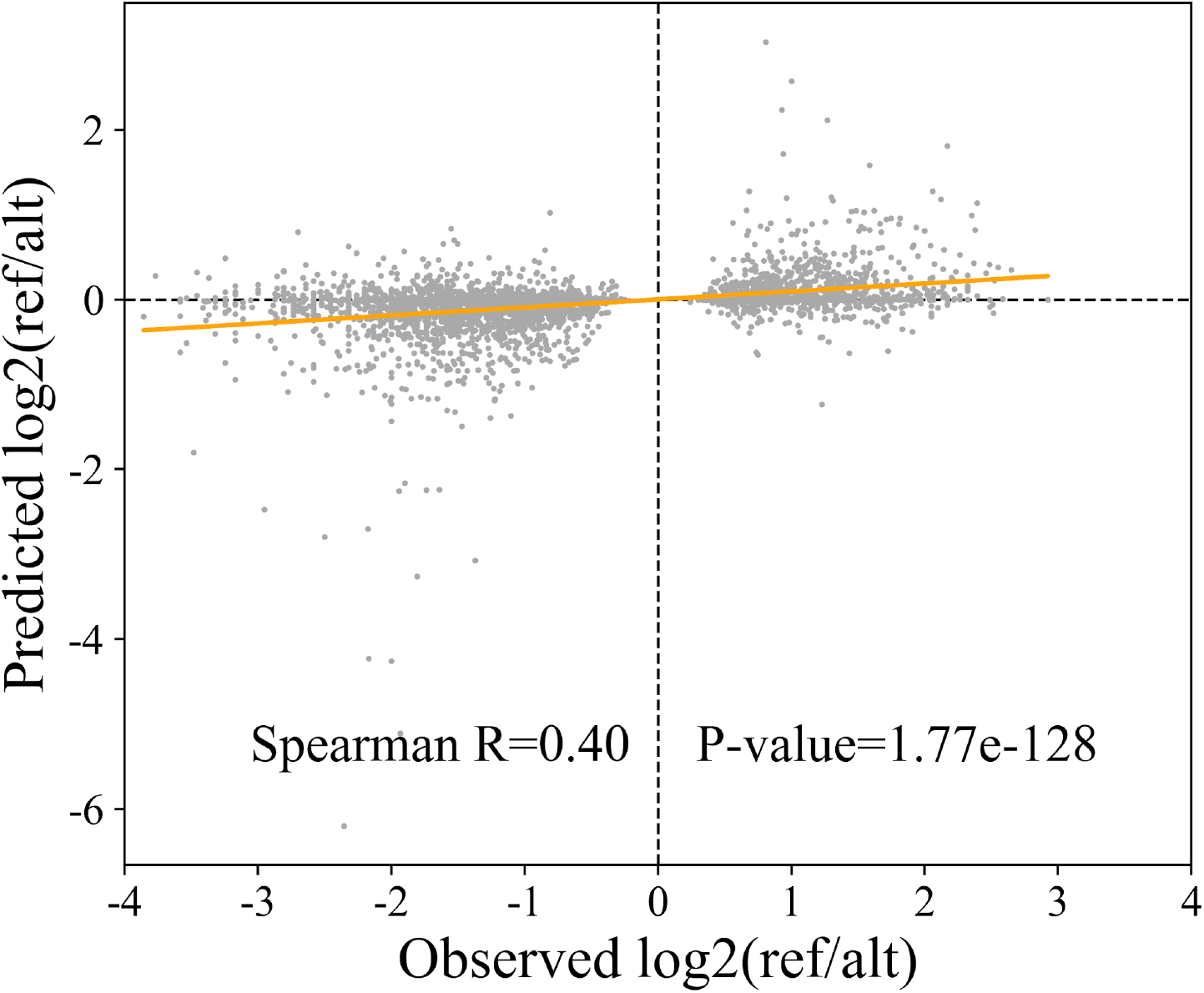
The observed allelic imbalance vs. MetaChrom predicted effects on chromatin accessibility of ASC variants in NPC neurons.

**Fig. S11:**
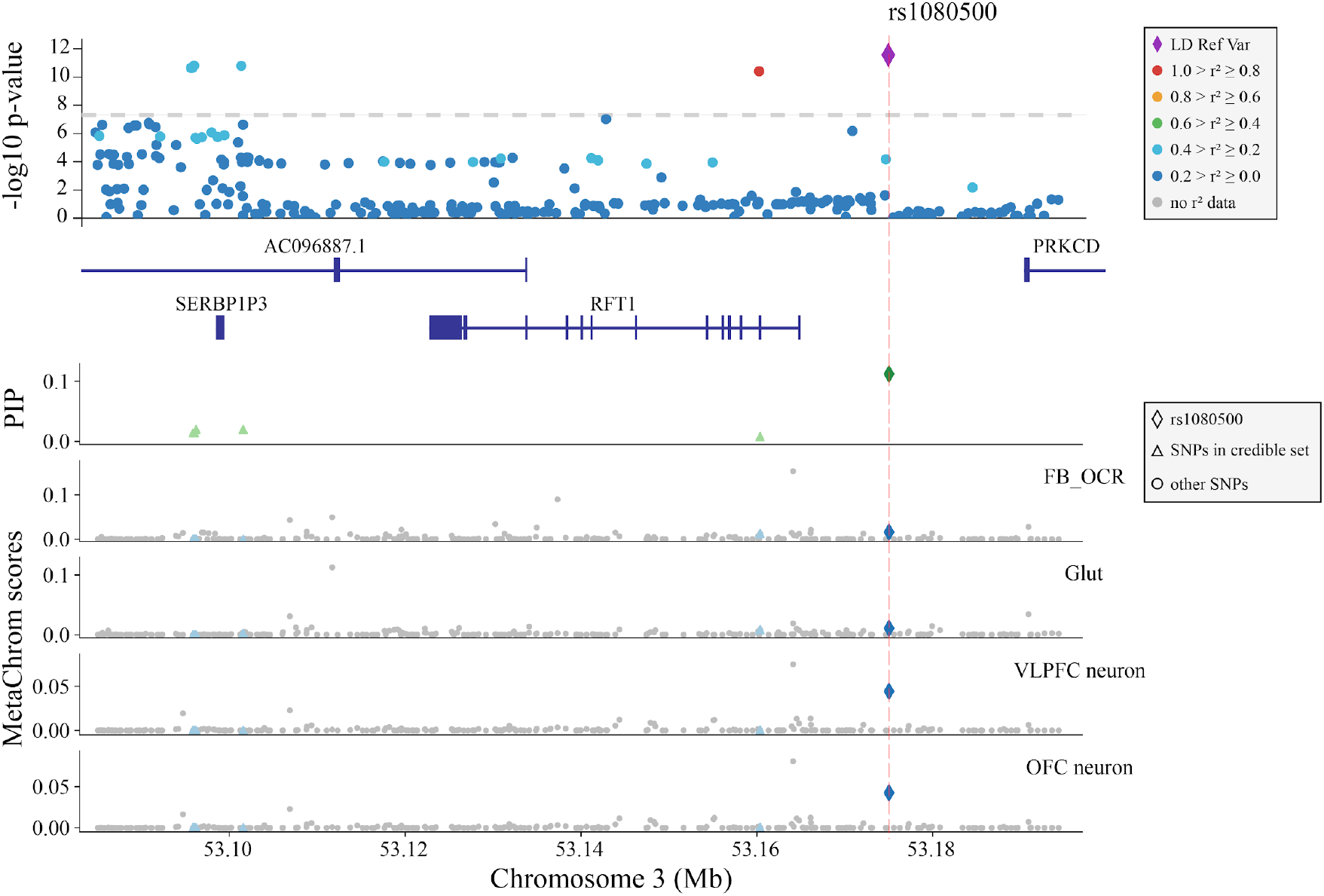
Likely causal variant rs1080500 and its MetaChrom functional annotations. The candidate SNP rs1080500 is chosen as the reference variant for computing LD and it is highlighted by the red dash line in each panel. The upper panel shows the significance of GWAS SNPs, LD between SNPs and genes in this region. The next panel shows credible set SNPs identified by fine-mapping (PIPs) in this region. The remaining panels show MetaChrom scores in four cell types, two in the fetal stage (FB_OCR and Glut) and two in the adult stage (VLPFC neuron and OFC neuron).

## References

1. B. Alipanahi, A. Delong, M. T. Weirauch, and B. J. Frey. Predicting the sequence specificities of dna-and rna-binding proteins by deep learning. Nature biotechnology, 33(8):831, 2015.

2. P. Arnold, A. Schöler, M. Pachkov, P. J. Balwierz, H. Jørgensen, M. B. Stadler, E. van Nimwegen, and D. Schübeler. Modeling of epigenome dynamics identifies transcription factors that mediate polycomb targeting. Genome research, 23(1):60–73, 2013.

3. Z. Avsec, R. Kreuzhuber, J. Israeli, N. Xu, J. Cheng, A. Shrikumar, A. Banerjee, D. S. Kim, L. Urban, A. Kundaje, et al. Kipoi: accelerating the community exchange and reuse of predictive models for genomics. BioRxiv, page 375345, 2018.

4. D. Benveniste, H.-J. Sonntag, G. Sanguinetti, and D. Sproul. Transcription factor binding predicts histone modifications in human cell lines. Proceedings of the National Academy of Sciences, 111(37):13367–13372, 2014.

5. B. E. Bernstein, J. A. Stamatoyannopoulos, J. F. Costello, B. Ren, A. Milosavljevic, A. Meissner, M. Kellis, M. A. Marra, A. L. Beaudet, J. R. Ecker, et al. The nih roadmap epigenomics mapping consortium. Nature biotechnology, 28(10):1045, 2010.

6. A.J. Bryois, M. E. Garrett, L. Song, A. Safi, P. Giusti-Rodriguez, G. D. Johnson, A. W. Shieh, Buil, J. F. Fullard, P. Roussos, et al. Evaluation of chromatin accessibility in prefrontal cortex of individuals with schizophrenia. Nature communications, 9(1):3121, 2018.

7. J. D. Buenrostro, B. Wu, H. Y. Chang, and W. J. Greenleaf. Atac-seq: a method for assaying chromatin accessibility genome-wide. Current protocols in molecular biology, 109(1):21–29, 2015.

8. D. Calderon, M. L. T. Nguyen, A. Mezger, A. Kathiria, F. Mller, V. Nguyen, N. Lescano, B. Wu, J. Trombetta, J. V. Ribado, et al. Landscape of stimulation-responsive chromatin across diverse human immune cells. Nature Genetics, 51(10):1494–1505, 2019.

9. J. Carlson, A. E. Locke, M. Flickinger, M. Zawistowski, S. Levy, R. M. Myers, M. Boehnke, H. M. Kang, L. J. Scott, J. Z. Li, et al. Extremely rare variants reveal patterns of germline mutation rate heterogeneity in humans. Nature communications, 9(1):1–13, 2018.

10. T. Ching, D. S. Himmelstein, B. K. Beaulieu-Jones, A. A. Kalinin, B. T. Do, G. P. Way, E. Ferrero, P.-M. Agapow, M. Zietz, M. M. Hoffman, et al. Opportunities and obstacles for deep learning in biology and medicine. Journal of The Royal Society Interface, 15(141):20170387, 2018.

11. G. P. Consortium et al. A global reference for human genetic variation. Nature, 526(7571):68, 2015.

12. E. P. Consortium et al. An integrated encyclopedia of dna elements in the human genome. Nature, 489(7414):57, 2012.

13. G. Consortium. Genetic effects on gene expression across human tissues. Nature, 550(7675):204–213, 2017.

14. R. Das, N. Dimitrova, Z. Xuan, R. A. Rollins, F. Haghighi, J. R. Edwards, J. Ju, T. H. Bestor, and M. Q. Zhang. Computational prediction of methylation status in human genomic sequences. Proceedings of the National Academy of Sciences, 103(28):10713–10716, 2006.

15. E. Davydov, D. L. Goode, M. Sirota, G. M. Cooper, A. Sidow, and S. Batzoglou. Identifying a high fraction of the human genome to be under selective constraint using gerp++. PLoS Computational Biology, 6(12):e1001025, 2010.

16. L. de la Torre-Ubieta, J. L. Stein, H. Won, C. K. Opland, D. Liang, D. Lu, and D. H. Geschwind. The dynamic landscape of open chromatin during human cortical neurogenesis. Cell, 172(1-2):289–304, 2018.

17. L. [de la Torre-Ubieta], J. L. Stein, H. Won, C. K. Opland, D. Liang, D. Lu, and D. H. Geschwind. The dynamic landscape of open chromatin during human cortical neurogenesis. Cell, 172(1):289 – 304.e18, 2018.

18. G. Eraslan, Ž. Avsec, J. Gagneur, and F. J. Theis. Deep learning: new computational modelling techniques for genomics. Nature Reviews Genetics, 20(7):389–403, 2019.

19. M. P. Forrest, H. Zhang, W. Moy, H. McGowan, C. Leites, L. E. Dionisio, Z. Xu, J. Shi, A. R. Sanders, W. J. Greenleaf, et al. Open chromatin profiling in hipsc-derived neurons prioritizes functional noncoding psychiatric risk variants and highlights neurodevelopmental loci. Cell stem cell, 21(3):305–318, 2017.

20. M. Fromer, P. Roussos, S. K. Sieberts, J. Johnson, D. H. Kavanagh, T. M. Perumal, D. M. Ruderfer, E. C. Oh, A. Topol, H. R. Shah, et al. Gene expression elucidates functional impact of polygenic risk for schizophrenia. Nature Neuroscience, 19(11):1442–1453, 2016.

21. J. F. Fullard, M. E. Hauberg, J. Bendl, G. Egervari, M.-D. Cirnaru, S. M. Reach, J. Motl, M. E. Ehrlich, Y. L. Hurd, and P. Roussos. An atlas of chromatin accessibility in the adult human brain. Genome research, 28(8):1243–1252, 2018.

22. J. F. Fullard, M. E. Hauberg, J. Bendl, G. Egervari, M.-D. Cirnaru, S. M. Reach, J. Motl, M. E. Ehrlich, Y. L. Hurd, and P. Roussos. An atlas of chromatin accessibility in the adult human brain. Genome research, 28(8):12431252, August 2018.

23. A. Garca-Bea, M. A. Walker, T. M. Hyde, J. E. Kleinman, P. J. Harrison, and T. A. Lane. Metabotropic glutamate receptor 3 (mglu3; mglur3; grm3) in schizophrenia: Antibody characterisation and a semi-quantitative western blot study. Schizophrenia research, 177(1-3):18–27, 2016.

24. R. E. Gate, C. S. Cheng, A. P. Aiden, A. Siba, M. Tabaka, D. Lituiev, I. Machol, M. G. Gordon, M. Subramaniam, M. Shamin, et al. Genetic determinants of co-accessible chromatin regions in activated t cells across humans. Nature Genetics, 50(8):1140–1150, 2018.

25. M. Genestine, L. Lin, M. Durens, Y. Yan, Y. Jiang, S. Prem, K. Bailoor, B. Kelly, P. K. Sonsalla, P. G. Matteson, et al. Engrailed-2 (en2) deletion produces multiple neurodevelopmental defects in monoamine systems, forebrain structures and neurogenesis and behavior. Human molecular genetics, 24(20):5805–5827, 2015.

26. D. L. Goode, G. M. Cooper, J. Schmutz, M. Dickson, E. Gonzales, M. Tsai, K. Karra, E. Davydov, S. Batzoglou, R. M. Myers, and A. Sidow. Evolutionary constraint facilitates interpretation of genetic variation in resequenced human genomes. Genome Research, 20(3):301, 2010.

27. A. G. B. Grønning, T. K. Doktor, S. J. Larsen, U. S. S. Petersen, L. L. Holm, G. H. Bruun, M. B. Hansen, A.-M. Hartung, J. Baumbach, and B. S. Andresen. Deepclip: predicting the effect of mutations on protein–rna binding with deep learning. Nucleic acids research, 48(13):7099–7118, 2020.

28. B. Gulko, M. J. Hubisz, I. Gronau, and A. Siepel. A method for calculating probabilities of fitness consequences for point mutations across the human genome. Nature genetics, 47(3):276, 2015.

29. S. Gupta, J. A. Stamatoyannopoulos, T. L. Bailey, and W. S. Noble. Quantifying similarity between motifs. Genome biology, 8(2):R24, 2007.

30. K. He, X. Zhang, S. Ren, and J. Sun. Deep residual learning for image recognition. In Proceedings of the IEEE conference on computer vision and pattern recognition, pages 770–778, 2016.

31. K. J. Karczewski, L. C. Francioli, G. Tiao, B. B. Cummings, J. Alfldi, Q. Wang, R. L. Collins, K. M. Laricchia, A. Ganna, D. P. Birnbaum, et al. The mutational constraint spectrum quantified from variation in 141,456 humans. Nature, 581(7809):434–443, 2020.

32. D. R. Kelley, J. Snoek, and J. L. Rinn. Basset: learning the regulatory code of the accessible genome with deep convolutional neural networks. Genome research, 26(7):990–999, 2016.

33. D. P. Kingma and J. Ba. Adam: A method for stochastic optimization. arXiv preprint arXiv:1412.6980, 2014.

34. P. K. Koo, P. Anand, S. B. Paul, and S. R. Eddy. Inferring sequence-structure preferences of rna-binding proteins with convolutional residual networks. bioRxiv, page 418459, 2018.

35. P. K. Koo and S. R. Eddy. Representation learning of genomic sequence motifs with convolutional neural networks. BioRxiv, page 362756, 2018.

36. A. Krizhevsky, I. Sutskever, and G. E. Hinton. Imagenet classification with deep convolutional neural networks. In Advances in neural information processing systems, pages 1097–1105, 2012.

37. A. Kundaje, W. Meuleman, J. Ernst, M. Bilenky, A. Yen, A. Heravi-Moussavi, P. Kheradpour, Z. Zhang, J. Wang, M. J. Ziller, et al. Integrative analysis of 111 reference human epigenomes. Nature, 518(7539):317, 2015.

38. J. H. Lam, Y. Li, L. Zhu, R. Umarov, H. Jiang, A. Héliou, F. K. Sheong, T. Liu, Y. Long, Y. Li, et al. A deep learning framework to predict binding preference of rna constituents on protein surface. Nature communications, 10(1):1–13, 2019.

39. D. Lee, D. U. Gorkin, M. Baker, B. J. Strober, A. L. Asoni, A. S. McCallion, and M. A. Beer. A method to predict the impact of regulatory variants from dna sequence. Nature genetics, 47(8):955, 2015.

40. M. Lek, K. J. Karczewski, E. V. Minikel, K. E. Samocha, E. Banks, T. Fennell, A. H. O’Donnell-Luria, J. S. Ware, A. J. Hill, B. B. Cummings, et al. Analysis of protein-coding genetic variation in 60,706 humans. Nature, 536(7616):258–291, 2016.

41. R. Leslie, C. J. ODonnell, and A. D. Johnson. Grasp: analysis of genotype-phenotype results from 1390 genome-wide association studies and corresponding open access database. Bioinformatics, 30(12):i185–i194, 2014.

42. C. Li and N. M. Luscombe. Nucleosome positioning stability is a modulator of germline mutation rate variation across the human genome. Nature communications, 11(1):1–13, 2020.

43. Z. Li and D. Hoiem. Learning without forgetting. IEEE transactions on pattern analysis and machine intelligence, 40(12):2935–2947, 2017.

44. Y. Liao, X. Zhuang, X. Huang, Y. Peng, X. Ma, Z.-X. Huang, F. Liu, J. Xu, Y. Wang, W.-M. Chen, et al. A bivalent securinine compound sn3-l6 induces neuronal differentiation via translational upregulation of neurogenic transcription factors. Frontiers in pharmacology, 9:290, 2018.

45. W. Liu, H. Zhou, L. Liu, C. Zhao, Y. Deng, L. Chen, L. Wu, N. Mandrycky, C. T. McNabb, Y. Peng, et al. Disruption of neurogenesis and cortical development in transgenic mice misexpressing olig2, a gene in the down syndrome critical region. Neurobiology of disease, 77:106–116, 2015.

46. L. Loewe. Negative selection. Nature Education, 1(1):59, 2008.

47. M. T. Maurano, R. Humbert, E. Rynes, R. E. Thurman, E. Haugen, H. Wang, A. P. Reynolds, R. Sandstrom, H. Qu, J. Brody, et al. Systematic localization of common disease-associated variation in regulatory dna. Science, 337(6099):1190–1195, 2012.

48. A. Melnik, S. Tauber, C. Dumrese, O. Ullrich, and S. Wolf. Murine adult neural progenitor cells alter their proliferative behavior and gene expression after the activation of toll-like-receptor 3. European journal of microbiology immunology, 2(3):239–248, 2012.

49. S. Nair, D. S. Kim, J. Perricone, and A. Kundaje. Integrating regulatory dna sequence and gene expression to predict genome-wide chromatin accessibility across cellular contexts. bioRxiv, page 605717, 2019.

50. V. Nair and G. E. Hinton. Rectified linear units improve restricted boltzmann machines. In Proceedings of the 27th international conference on machine learning (ICML-10), pages 807–814, 2010.

51. Y. E. Nesterov. A method for solving the convex programming problem with convergence rate o (1/k^ 2). In Dokl. akad. nauk Sssr, volume 269, pages 543–547, 1983.

52. S. W. G. of the Psychiatric Genomics Consortium et al. Biological insights from 108 schizophrenia-associated genetic loci. Nature, 511(7510):421–427, 2014.

53. C.-T. Ong and V. G. Corces. Ctcf: an architectural protein bridging genome topology and function. Nature reviews Genetics, 15(4):234, 2014.

54. R. H. Paap, S. Oosterbroek, C. M. Wagemans, L. von Oerthel, R. D. Schellevis, A. J. Vastenhouw-van der Linden, M. J. G. Koerkamp, M. F. Hoekman, and M. P. Smidt. Foxo6 affects plxna4-mediated neuronal migration during mouse cortical development. Proceedings of the National Academy of Sciences, 113(45):E7087–E7096, 2016.

55. J. T. M. L. Paridaen, E. Janson, K. H. Utami, T. C. Pereboom, P. B. Essers, C. v. Rooijen, D. Zivkovic, and A. W. Maclnnes. The nucleolar gtp-binding proteins gnl2 and nucleostemin are required for retinal neurogenesis in developing zebrafish. Developmental biology, 355(2):286–301, 2011.

56. P. J. Park. Chip–seq: advantages and challenges of a maturing technology. Nature reviews genetics, 10(10):669, 2009.

57. J. K. Pickrell. Joint analysis of functional genomic data and genome-wide association studies of 18 human traits. The American Journal of Human Genetics, 94(4):559–573, 2014.

58. L. Pinello, J. Xu, S. H. Orkin, and G.-C. Yuan. Analysis of chromatin-state plasticity identifies cell-type–specific regulators of h3k27me3 patterns. Proceedings of the National Academy of Sciences, 111(3):E344–E353, 2014.

59. D. Quang and X. Xie. Danq: a hybrid convolutional and recurrent deep neural network for quantifying the function of dna sequences. Nucleic acids research, 44(11):e107–e107, 2016.

60. S. K. Reilly, J. Yin, A. E. Ayoub, D. Emera, J. Leng, J. Cotney, R. Sarro, P. Rakic, and J. P. Noonan. Evolutionary changes in promoter and enhancer activity during human corticogenesis. 347(6226):1155–1159, 2015.

61. P. Rentzsch, D. Witten, G. M. Cooper, J. Shendure, and M. Kircher. Cadd: predicting the deleteriousness of variants throughout the human genome. Nucleic acids research, 47(D1):D886–D894, 2018.

62. R. Sabarinathan, L. Mularoni, J. Deu-Pons, A. Gonzalez-Perez, and N. López-Bigas. Nucleotide excision repair is impaired by binding of transcription factors to dna. Nature, 532(7598):264–267, 2016.

63. J. Schreiber, R. Singh, J. Bilmes, and W. S. Noble. A pitfall for machine learning methods aiming to predict across cell types. Genome biology, 21(1):1–6, 2020.

64. N. Schrode, S.-M. Ho, K. Yamamuro, A. Dobbyn, L. Huckins, M. R. Matos, E. Cheng, P. J. M. Deans, E. Flaherty, N. Barretto, et al. Synergistic effects of common schizophrenia risk variants. Nature genetics, 51(10):1475–1485, 2019.

65. M. Setty and C. S. Leslie. Seqgl identifies context-dependent binding signals in genome-wide regulatory element maps. PLoS computational biology, 11(5):e1004271, 2015.

66. A. Shrikumar, P. Greenside, and A. Kundaje. Learning important features through propagating activation differences. In Proceedings of the 34th International Conference on Machine Learning-Volume 70, pages 3145–3153. JMLR. org, 2017.

67. A. Shrikumar, K. Tian, A. Shcherbina, Ž. Avsec, A. Banerjee, M. Sharmin, S. Nair, and A. Kundaje. Tf-modisco v0. 4.4. 2-alpha. arXiv preprint arXiv:1811.00416, 2018.

68. K. Simonyan, A. Vedaldi, and A. Zisserman. Deep inside convolutional networks: Visualising image classification models and saliency maps. arXiv preprint arXiv:1312.6034, 2013.

69. R. Socher, Y. Bengio, and C. D. Manning. Deep learning for nlp (without magic). In Tutorial Abstracts of ACL 2012, pages 5–5. Association for Computational Linguistics, 2012.

70. L. Song and G. E. Crawford. Dnase-seq: a high-resolution technique for mapping active gene regulatory elements across the genome from mammalian cells. Cold Spring Harbor Protocols, 2010(2):pdb–prot5384, 2010.

71. M. Song, X. Yang, X. Ren, L. Maliskova, B. Li, I. R. Jones, C. Wang, F. Jacob, K. Wu, M. Traglia, et al. Mapping cis-regulatory chromatin contacts in neural cells links neuropsychiatric disorder risk variants to target genes. Nature Genetics, 51(8):1252–1262, 2019.

72. Z. Sun, C. S. da Fontoura, M. Moreno, N. E. Holton, M. Sweat, Y. Sweat, M. K. Lee, J. Arbon, F. B. Bidlack, D. R. Thedens, et al. Foxo6 regulates hippo signaling and growth of the craniofacial complex. PLoS genetics, 14(10):e1007675, 2018.

73. K. Takahashi and S. Yamanaka. Induction of pluripotent stem cells from mouse embryonic and adult fibroblast cultures by defined factors. cell, 126(4):663–676, 2006.

74. C. Tan, F. Sun, T. Kong, W. Zhang, C. Yang, and C. Liu. A survey on deep transfer learning. In International Conference on Artificial Neural Networks, pages 270–279. Springer, 2018.

75. S. B. Thyme, L. M. Pieper, E. H. Li, S. Pandey, Y. Wang, N. S. Morris, C. Sha, J. W. Choi, K. J. Herrera, E. R. Soucy, et al. Phenotypic landscape of schizophrenia-associated genes defines candidates and their shared functions. Cell, 177(2):478–491, 2019.

76. A. E. Trevino, N. Sinnott-Armstrong, J. Andersen, S.-J. Yoon, N. Huber, J. K. Pritchard, H. Y. Chang, W. J. Greenleaf, and S. P. Pasca. Chromatin accessibility dynamics in a model of human forebrain development. 367(6476), 2020.

77. M. Tsompana and M. J. Buck. Chromatin accessibility: a window into the genome. Epigenetics & chromatin, 7(1):1–16, 2014.

78. K. C. Vadodaria, D. N. Amatya, M. C. Marchetto, and F. H. Gage. Modeling psychiatric disorders using patient stem cell-derived neurons: a way forward. Genome medicine, 10(1):1, 2018.

79. F. N. Velazquez, B. L. Caputto, and F. D. Boussin. c-fos importance for brain development. Aging (Albany NY), 7(12):1028, 2015.

80. R. Walker, G. Ramaswami, C. Hartl, N. Mancuso, M. J. Gandal, L. d. l. Torre-Ubieta, B. Pasaniuc, J. L. Stein, and D. H. Geschwind. Genetic control of expression and splicing in developing human brain informs disease mechanisms. Cell, 179(3):750–771, 2019.

81. A. Wang, F. Yue, Y. Li, R. Xie, T. Harper, N. A. Patel, K. Muth, J. Palmer, Y. Qiu, J. Wang, et al. Epigenetic priming of enhancers predicts developmental competence of hesc-derived endodermal lineage intermediates. Cell stem cell, 16(4):386–399, 2015.

82. D. Wang, S. Liu, J. Warrell, H. Won, X. Shi, F. C. P. Navarro, D. Clarke, M. Gu, P. Emani, Y. T. Yang, M. Xu, M. J. Gandal, S. Lou, J. Zhang, J. J. Park, C. Yan, S. K. Rhie, K. Manakongtreecheep, H. Zhou, A. Nathan, M. Peters, E. Mattei, D. Fitzgerald, T. Brunetti, J. Moore, Y. Jiang, K. Girdhar, G. E. Hoffman, S. Kalayci, Z. H. Gümüş, G. E. Crawford, , P. Roussos, S. Akbarian, A. E. Jaffe, K. P. White, Z. Weng, N. Sestan, D. H. Geschwind, J. A. Knowles, and M. B. Gerstein. Comprehensive functional genomic resource and integrative model for the human brain. 362(6420), 2018.

83. k. Wang, M. Li, and H. Hakon. Annovar: Functional annotation of genetic variants from next-generation sequencing data. Nucleic acids research, 38(16):e164, 2010.

84. S. Wang, S. Sun, Z. Li, R. Zhang, and J. Xu. Accurate de novo prediction of protein contact map by ultra-deep learning model. PLoS computational biology, 13(1):e1005324, 2017.

85. M. T. Weirauch, A. Yang, M. Albu, A. G. Cote, A. Montenegro-Montero, P. Drewe, H. S. Najafabadi, S. A. Lambert, I. Mann, K. Cook, et al. Determination and inference of eukaryotic transcription factor sequence specificity. Cell, 158(6):1431–1443, 2014.

86. X. Wen, Y. Lee, F. Luca, and R. Pique-Regi. Efficient integrative multi-snp association analysis via deterministic approximation of posteriors. The American Journal of Human Genetics, 98(6):1114–1129, 2016.

87. J. W. Whitaker, Z. Chen, and W. Wang. Predicting the human epigenome from dna motifs. Nature methods, 12(3):265, 2014.

88. b. Xiang, Q. Wang, W. Lei, M. Li, Y. Li, L. Zhao, X. Ma, Y. Wang, H. Yu, X. Li, et al. Genes in immune pathways associated with abnormal white matter integrity in first-episode and treatment-nave patients with schizophrenia. The British journal of psychiatry, 214(5):281–287, 2019.

89. N. Yutsudo, T. Kamada, K. Kajitani, H. Nomaru, A. Katogi, Y. H. Ohnishi, Y. N. Ohnishi, K.-i. Takase, K. Sakumi, H. Shigeto, et al. fosb-null mice display impaired adult hippocampal neuro-genesis and spontaneous epilepsy with depressive behavior. Neuropsychopharmacology, 38(5):895, 2013.

90. S. Zhang, H. Zhang, Y. Zhou, M. Qiao, S. Zhao, A. Kozlova, J. Shi, A. R. Sanders, G. Wang, K. Luo, et al. Allele-specific open chromatin in human ipsc neurons elucidates functional disease variants. Science, 369(6503):561–565, 2020.

91. J. Zhou and O. G. Troyanskaya. Predicting effects of noncoding variants with deep learning–based sequence model. Nature methods, 12(10):931, 2015.

